# The representation of visual motion and landmark position aligns with heading direction in the zebrafish interpeduncular nucleus

**DOI:** 10.1101/2024.09.25.614953

**Authors:** Hagar Lavian, Ot Prat, Luigi Petrucco, Vilim Štih, Ruben Portugues

**Affiliations:** Institute of Neuroscience, Technical University of Munich, Germany; Graduate School of Systemic Neurosciences, Ludwig-Maximilian University, Munich, Germany; Center for Neuroscience and Cognitive Systems, Istituto Italiano di Tecnologia, Rovereto, Italy; EnliteAI, Vienna, Austria; Bernstein Center for Computational Neuroscience, Munich, Germany; SyNergy Excellence Cluster, Munich, Germany; Max Planck Fellow Group - Mechanisms of Cognition, MPI Psychiatry, Munich, Germany

## Abstract

Sensory information is fundamental for navigation. Visual motion is used by animals to estimate their traveling distance and direction, and visual landmarks allow animals to tether their location and orientation to their environment. How such signals are integrated in the vertebrate brain is poorly understood. Here we investigate the representation of directional whole field visual motion and landmark position in a circuit in the larval zebrafish consisting of the habenula, interpeduncular nucleus (IPN) and anterior hindbrain (aHB). This circuit has been recently implicated in the representation of heading direction. Using calcium imaging we show that these stimuli are represented in the habenula, IPN and aHB. We further show that their representation in the IPN of both these stimuli is topographically arranged in a way that aligns itself with the representation of the heading signal in this region. We use neuronal ablations to show that the landmark responses, but not the whole field motion responses, require intact habenula input to the IPN. Overall our findings suggest the IPN as a site for integration of the heading signal from the aHB with visual information, shedding light on how different types of navigational signals are processed in the vertebrate brain.

## Introduction

To navigate the environment, an animal must know where it is heading. Neural networks that represent the animal’s orientation within its environment, called heading direction (HD) networks, have been found in mammals, birds, fish and insects [1–6]. In the absence of landmarks, animals can path integrate self-motion cues such as vestibular input, self-generated optic flow and motor commands to maintain an updated internal representation of heading [7–13]. This path integration is nevertheless prone to the accumulation of errors and HD networks are less stable in the absence of visual landmarks [14, 15]. Anchoring the HD network therefore requires the incorporation of sensory input into this internal representation, but the mechanisms by which this is achieved in vertebrates remain largely unknown.

In mammals, HD neurons (HDNs) have been found in several brain regions, including the dorsal tegmental nucleus (DTN), lateral mammillary nucleus (LMN), anterodorsal thalamic nucleus (ADN), postsubiculum (Po-Sub) and entorhinal and retrosplenial cortices [7, 13, 16–19]. The heading signal is internally generated by a network that integrates vestibular signals and motor commands, most likely in the DTN and LMN [18, 20–23]. This signal is transmitted to the thalamus and PoSub. Previous studies have shown that activity of HDNs in the ADN, PoSub and retrosplenial cortex is sensitive to visual information [1, 14, 24]. While some studies suggest that sensory information is most likely integrated in the Po-Sub [25] or the retrosplenial cortex [24, 26], it is still not known whether such information affects HDNs in other parts of the network.

In recent years, the topography and connectivity of the insect heading direction network has been discovered and described in great detail [5, 27–31]. The insect HDNs form a network in a structure called the central complex in the center of the insect brain [5]. The neuropil of the insect HDNs is organized as wedges in the ellipsoid body, a circular structure located within the central complex. Heading direction representation in this network can be shifted by motor information (if the animal turns) or by sensory information, such as visual or somatosensory cues (e.g., if a landmark in the environment moves to a new location) [5, 32–36]. This rotation is performed by shift neurons, which have their axons tiling the ellipsoid body in a similar manner to the HDNs but with a small offset. Surprisingly, the neurons that provide sensory information to this system do not form localized arborizations but rather connect to all HDNs [31, 37]. The fact that many other types of neurons that innervate the ellipsoid body, or other parts of the central complex, tile a specific wedge, row or column in this region [31, 38] allows for local modulation of these global inputs. This principle of organization appears to allow for very flexible routes of information flow in the insect navigation system.

The insect central complex is also a site of integration of different signals that are relevant for navigation, such as heading and the animal’s traveling direction. While HDNs themselves are not sensitive to translational visual motion, another type of cell in this region responds to visual motion in a particular direction [39, 40]. It is currently unclear where such integration exists in the vertebrate brain.

Recently, we discovered a heading direction network in the zebrafish anterior hindbrain (aHB) and suggested this as the homologous structure of the mammalian DTN [6]. This is due to three reasons: (1) the DTN develops from rhombomere 1, where the zebrafish HDN is located, (2) both regions contain a group of GABAergic HD neurons and (3) the aHB and DTN show similar connectivity patterns with the habenula and interpeduncular nucleus (IPN) [13, 18, 41–43]. Here, we investigate the representation of different types of visual information in the aHB-IPN-habenula network. We show that all three structures contain visually tuned neurons. We further show that direction of optic flow and landmark position are topographically represented in the dorsal IPN (dIPN). Finally, we use ablations to show that habenula input is required for representation of landmark position in the dIPN, but not for representation of visual motion or basic functioning of the HD network.

## Results

### Neurons in the left habenula, aHB and IPN are tuned to directional visual motion

Our first goal was to detect neurons, throughout the brain, tuned to directional visual motion, a visual signal that indicates an animal’s traveling direction. We used lightsheet microscopy to obtain whole brain imaging datasets from Tg(Huc:H2B-GCaMP6s) zebrafish larvae presented with whole field translational visual motion (Figure 1A) (see Methods). In each trial, the projected pattern moved in one of the eight cardinal or intercardinal directions with respect to the fish, chosen in a random order. Neurons responding reliably to visual motion in at least one direction were detected in the tectum, pretectum, left habenula, aHB and IPN, in line with previous studies [44–46] (Figure 1B-F, Figure S1-Figure S3). Out of the regions containing reliably responding neurons (see Methods for reliability index calculation), the representation of different directions in the pretectum and aHB was found to be lateralized such that neurons responding to leftward motion are on the left hemisphere of the brain, and neurons responding to rightward motion are on the right hemisphere. In contrast, neurons in the left habenula and tectum do not exhibit this lateralization and visually tuned neurons in these regions do not have a particular anatomical distribution (Figure 1C-D).

**Figure 1:**
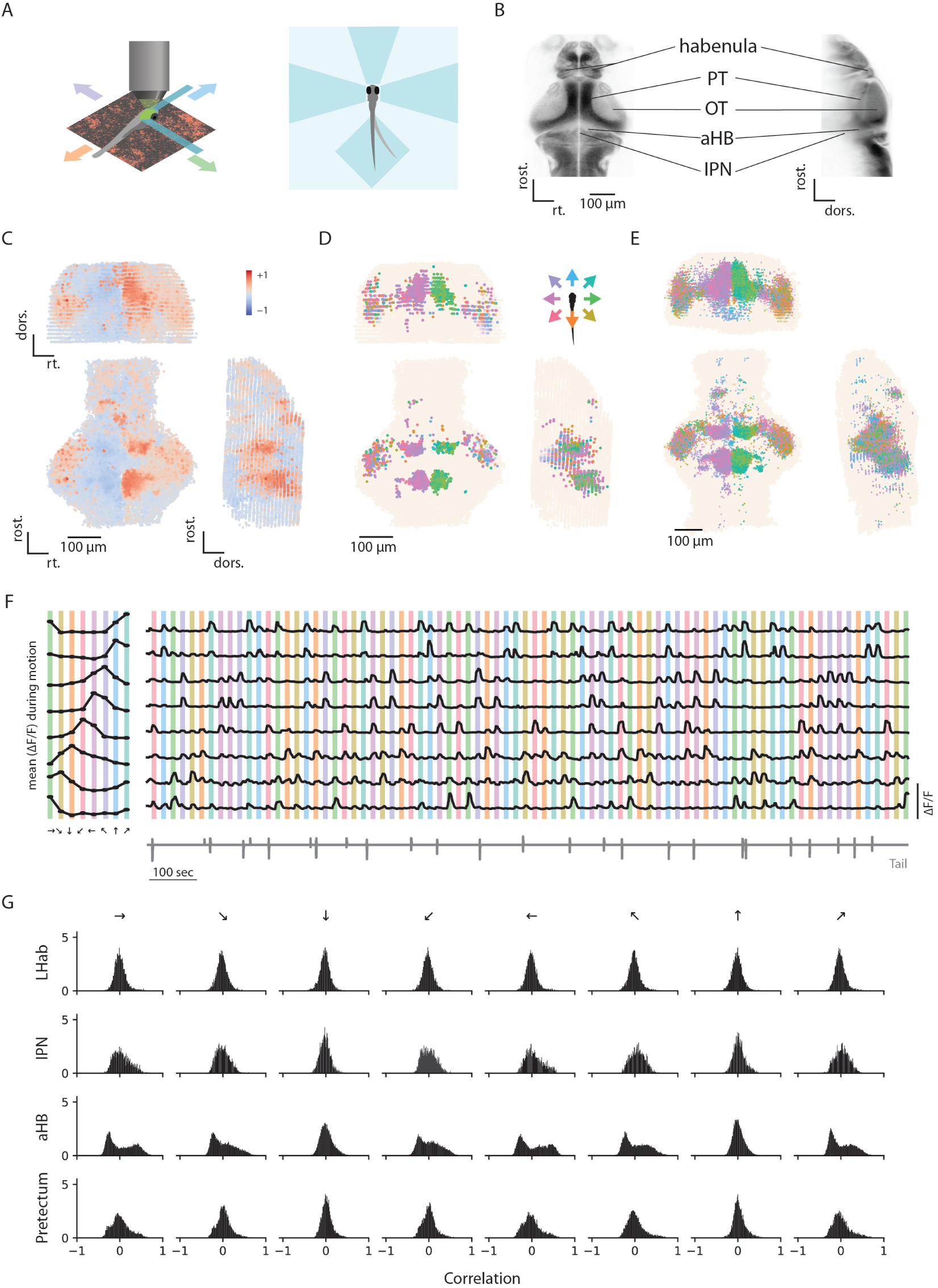
Whole brain responses to whole field visual motion. (A) Illustration of experimental design. Left, an embedded zebrafish on top of a pink noise pattern moving in 8 possible directions. Right, embedded zebrafish larvae in a lightsheet imaging chamber. Agarose is removed from the tail to allow swimming and tail tracking. Agarose is also removed from the side and front to prevent the laser from scattering. (B) horizontal and sagittal views of the zebrafish brain. Lines point to specific regions that were found to respond to whole field visual motion. PT: pretectum, OT: optic tectum, aHB: anterior hindbrain, IPN: interpeduncular nucleus. (C) Three views of all ROIs extracted from a whole brain dataset (one example fish). Neurons are colored according to their correlation value with rightward motion. (D) Three views of the same fish, only reliably responding neurons are shown (top 5% of reliability index). Each neuron is colored according to the direction it is tuned to. (E) Same as D but for all fish in the dataset, registered to a reference fish (n=15). (F) Example neurons that are tuned to different directions. Left, tuning curves of 8 neurons. Right, the full traces from the entire experiment for the same neurons. The colorful shadings indicate the direction that is being presented. (G) Distributions of correlation values with each direction for neurons in the left habenula (Lhab), IPN, aHB and pretectum (n=15).

### Head direction neurons and visually tuned neurons are separate populations in the aHB

The aHB, the region containing the zebrafish HDNs, also contains many neurons that are tuned to visual motion (Figure 1). We next wanted to check if HDNs in the aHB respond to visual motion. We used a lightsheet microscope to image GABAergic neurons in the aHB in both darkness and while presenting directional whole field motion (Figure 2A). HDNs were selected based on their activity in the darkness phase as previously described in [6]. Our results showed that HDNs and visually tuned neurons form two non-overlapping populations (Figure 2). In all of our fish, neurons classified as HDNs did not respond to translational visual motion (Figure 2C-D). Similarly, neurons that reliably respond to the visual stimulus were never identified as HDNs, i.e. they were not negatively correlated with other neurons during the darkness phase (Figure 2C-D).

**Figure 2:**
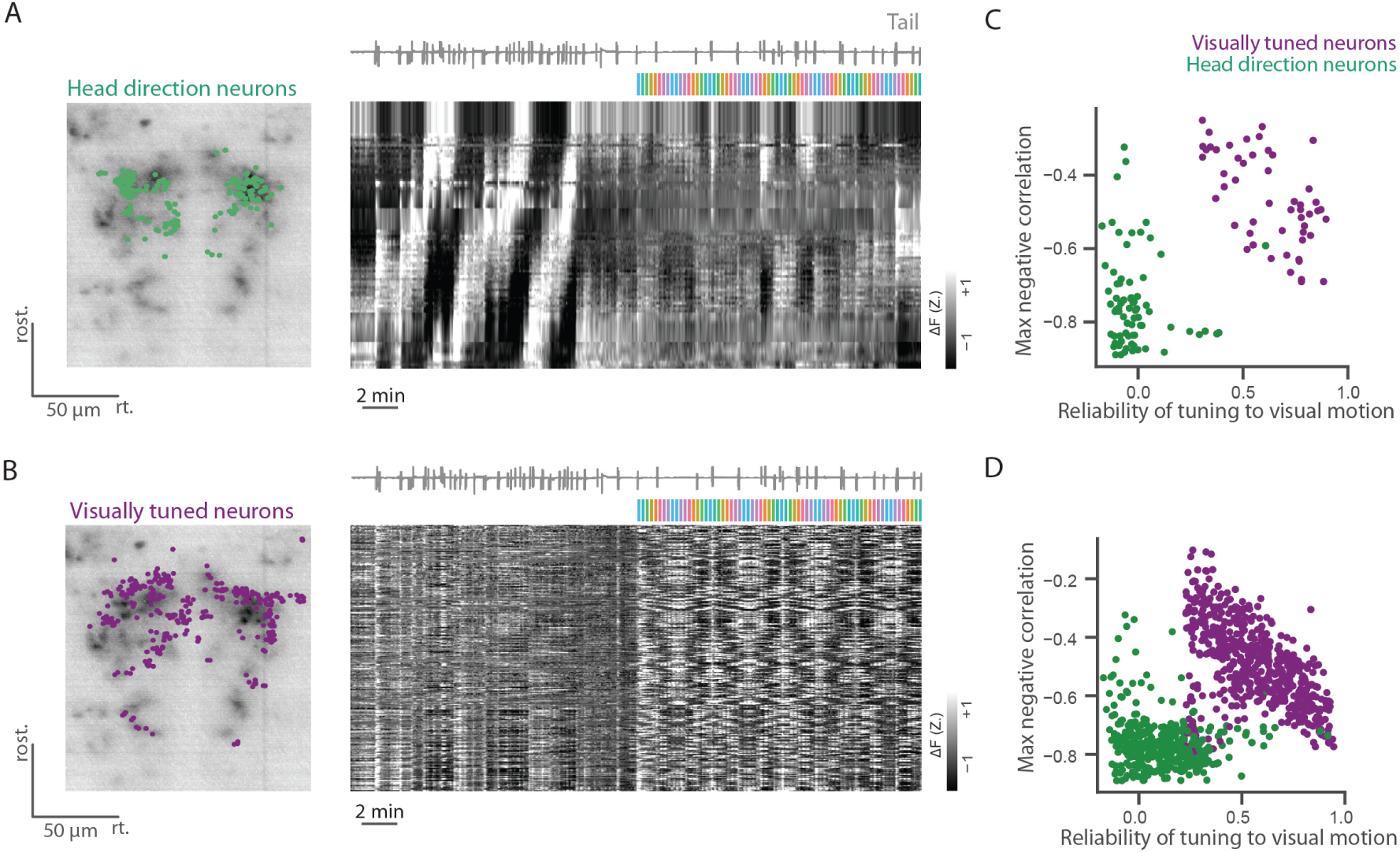
Visually tuned neurons and heading direction neurons are two separate populations in the aHB. (A) Left, anatomy of the aHB of a Tg(gad1b:Gal4; UAS:GCaMP6s) fish. HDNs are indicated in green. Right, traces of heading direction neurons, sorted according to their phase. The tail trace and colorful shading indicating presentation of visual motion are shown on top of the traces. (B) Left, anatomy of the aHB in the same fish. Visually tuned neurons are labeled in purple. Right, traces of visually tuned neurons. (C) Scatter plot showing the HDNs and the visually tuned neurons are functionally two separate populations (same example fish as in panels A and B). HDNs do not show reliable responses to visual motion (x axis) and show more negative correlation values with other neurons during the darkness phase of the experiment (y axis). Heading direction neurons have more negative correlation values with other neurons during darkness and visually tuned neurons have higher values of there liability index. (D) Sameas C but for all fish (n(fish)=3, n(HDNs)=[74, 172, 137], n(visually tuned)=[56, 367, 236]).

### Visual motion is topographically arranged in the dIPN

Our whole brain imaging data revealed neurons tuned to directional whole field motion in the IPN, where HDNs from the aHB project their dendrites and axons to [6]. We next imaged different transgenic lines that label different components of the IPN, and characterized their responses to directional visual motion. To characterize the tuning of IPN neuropil to visual motion, we generated tuning maps in which each pixel is colored according to the direction it is tuned to (Figure 3, Figure S4, Figure S5). We first imaged Tg(elavl3:GCaMP6s) fish, in which GCaMP6s is expressed in the cytosol of all neurons, labeling both somata and neuropil. In this pan-neuronal line, we noticed a strong tuning to visual motion in the dIPN. This activity is topographically organized, with each direction represented in a single column on the rostro-caudal axis. The leftmost column is tuned to backward-left motion and the rightmost column is tuned to backward-right motion (Figure 3C-D). This topographic organization is highly consistent across different planes in the dIPN (Figure S4) and across fish (Figure 3). To better understand how this columnar structure is formed, we imaged fish lines with more restricted expression patterns. Next, we imaged the Tg(16715:Gal4; UAS:GCaMP6s) line, labeling only neurons in the habenula, a major source of excitatory input to the IPN in mammals and fish [47–50]. In zebrafish, the left habenula projects to both dorsal and ventral parts of the IPN while the right habenula projects mostly to the ventral IPN [51, 52]. It is further known that habenular neurons project their axons to the IPN, where they wrap around the structure [53]. Despite this global anatomical organization, activity of habenular axons in the dIPN shows a topographical tuning to the visual stimulus. Similarly to the pattern we observed in the Tg(elavl3:GCaMP6s) line (Figure 3C-D), habenula axons show tuning organized in columns in the rostro-caudal axis (Figure 3E-F). This organization is surprising because, as mentioned above, habenular neurons are known to largely wrap around the dIPN and they do not form localized projections.

**Figure 3:**
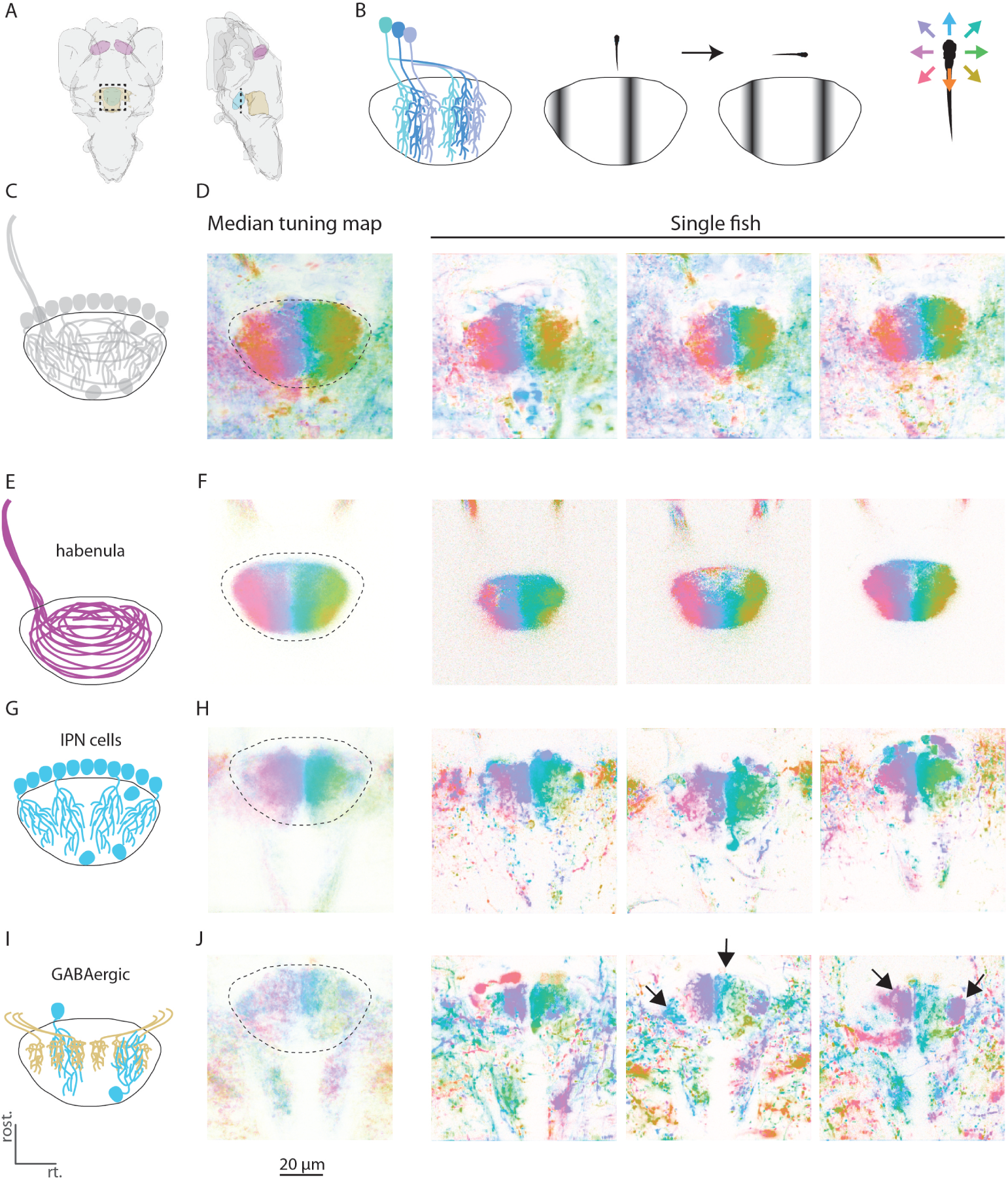
Representation of visual motion tuning in the dIPN is organized in columns. (A), Illustration of the zebrafish brain, horizontal (left) and sagittal (right) views. The IPN and its afferent structures are highlighted (habenula is shown in purple, IPN is shown in light blue, aHB is shown in gold) dashed square indicates the region images in following plots. (B) Illustration of heading direction representation in the dIPN (based on [6]). Left, illustration of the morphology of three HDNs and their projections in the dIPN: each HDN has a single neurite that goes ventrally and splits into an ipsilateral dendrite and a contralateral axon. Right, illustration of the activity of HDN neuropil in the IPN when the fish is facing different directions. When an HDN is active, both its axon and dendrite are active in the dIPN (C) Illustration of the IPN and its different components. (D) Left, median tuning map of the dIPN in the Tg(elavl3:GCaMP6s) line labeling the cytosol pan-neuronally (n=4). Each pixel is colored according to the direction it is tuned to. Right, tuning maps of three example fish. (E) Illustration of habenula input to the dIPN. Axons from the left habenula enter the IPN and then wrap around it. (F) Left, median tuning map of the dIPN in the Tg(16715:Gal4) line labeling only habenula axons (n=12). Right, tuning maps of three example fish. (G) Illustration of IPN cells and their neuropil. (H) Left, median tuning map of the dIPN in the Tg(s1168t:Gal4) line labeling IPN cells and neuropil (n=15). Right, tuning maps of three example fish. (I) Illustration of GABAergic elements in the dIPN, which consists mostly from sparse expression in the IPN itself with additional neuropil that mostly originates in the aHB. (J) Left, median tuning map of the dIPN in the Tg(gad1b:Gal4) fish (n=11). Right, tuning maps of three example fish. Arrows indicate neuropil columns that are tuned to the same direction on both sides of the dIPN.

We next imaged the Tg(s1168t:Gal4; UAS:GCaMP6s) fish line labeling IPN neurons and their neuropil, but none of the structures that project to the IPN (Figure S6). The cell bodies of these neurons are aligned from left to right along the rostral boundary of the dIPN, and their neuropil extends caudally. This neuropil is tuned to directional motion in a very similar manner to that found in the habenula axons (Figure 3G-H). This tuning is organized in a columnar structure, such that left columns are tuned to leftward visual motion and right columns are tuned to rightward visual motion. The overall organization is very similar to that observed in the pan-neuronal and habenula lines, with a single exception of lack of tuning to backward motion by IPN cells (Figure S5).

We next imaged Tg(gad1b:Gal4; UAS:GCaMP6s) fish, in which only GABAergic neurons are labeled (Figure 3I-J). When imaging the GABAergic neuropil in the dIPN, we imaged dendrites and axons of IPN neurons as well as those of aHB and potentially other GABAergic neurons projecting to the dIPN. These experiments revealed that the GABAergic neuropil in the dIPN is strongly tuned to visual motion in different directions. The tuning appears again to be organized in a columnar structure, such that different directions are represented by different neuropil columns (Figure 3I). However, the organization of the GABAergic neuropil is different from that observed for IPN cells and habenula axons. While in the previous datasets tuning to each direction appeared in a single column, when imaging the GABAergic neuropil we found that each direction appears on both sides of the IPN. This is very similar to the structure of the heading direction signal in the dIPN (Figure 3B, [6]). As our data shows that HDNs themselves do not respond to visual motion, in the likely case that this tuning arises from aHB neurons, this would suggest that visually tuned neurons in the aHB have a similar morphology to that of HDNs.

### Landmark position is topographically arranged in the dIPN

Many animals use visual landmarks to anchor their heading direction to their sensory environment. We next wanted to check if landmark position is at all represented in the dIPN. Neurons that have local receptive fields could represent this type of information. To detect such neurons, we used a two-photon microscope to record activity of IPN neurons in zebrafish larvae that were presented with a light bar at different azimuth angles in their front visual field (Figure 4A, see Methods).

**Figure 4:**
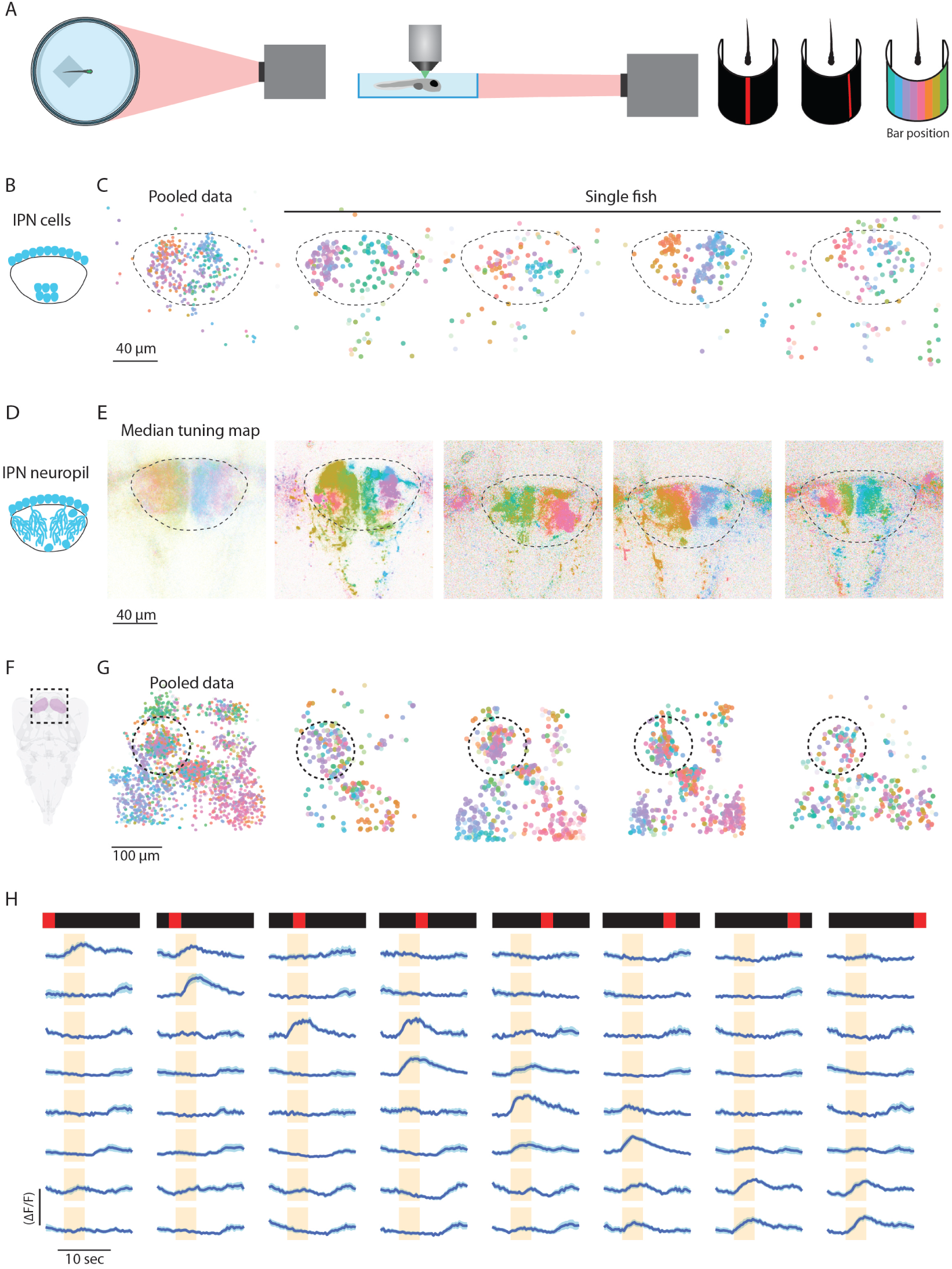
Representation of light position in the dIPN and the habenula. (A) Illustration of experimental setup of presentation of frontal visual stimulus to embedded fish. (B) Illustration of somata distribution in the dIPN as seen in the Tg(elavl3:H2B-GCaMP6s) line. (C) Tuning of dIPN cells to light position. Only reliably responding ROIs are shown, each ROI is colored according to the position it is tuned to. Left, data from multiple fish registered to one another (n=7). Right, same as in the left panel but for four individual fish. The dIPN in each dataset is marked with a dashed circle. (D) Illustration of dIPN somata and neuropil. (E) Tuning of dIPN neuropil to landmark position. Left, average tuning map of Tg(s1168t:Gal4; UAS:GCaMP6s) fish, showing the average response pattern, each pixel is colored according to the direction that it is tuned to (n=11). Right, example fish tuning maps of the dIPN in three example fish. (F) Illustration of the zebrafish brain, the habenula is highlighted in pink. (G) Tuning of habenula cells to landmark position as found in Tg(elavl3:H2B-GCaMP6s) and Tg(elavl3:GCaMP6s) fish. Only reliably responding ROIs are shown, each ROI is colored according to the position it is tuned to. Left, data from multiple fish morphed to one another (n=16). Right, same as in the left panel but for four individual fish. The left habenula in each dataset is marked with a dashed circle. (H) Responses of eight neurons from the left habenula to the appearance of a light bar in the eight possible locations. For each neuron the mean ± sem response is shown. Orange shading indicates stimulus presentation.

When imaging cell bodies in the IPN, we found neurons, in all fish, that responded to light in a particular position of the visual field. In all fish, these light-responsive neurons tiled the IPN retinotopically. In addition, for most fish, this retinotopic map was aligned: neurons on the right side of the IPN responded to light on the right side of the visual field, whereas neurons on the left responded to stimuli on the left (Figure 4B-C). For some fish though, (see first individual fish in 4C), this map was ”phase-shifted”, i.e. there was a constant offset between the left-right alignment of the visual field and the left-right alignment in the IPN.

We next wanted to better characterize the representation of landmark position in the dIPN. To do this we imaged the IPN neuropil while showing the same light bar stimulus. In these experiments, we observed a more graded pattern than we could observe in the somata, with the azimuthal angle in the visual field similarly encoded in the left-right dIPN axis in columns that extended rostro-caudally (Figure 4D-E), similar to that observed in response to visual motion (Figure 3G) and to the neuropil of HDNs (Figure 3B and [6]). The columnar structure is consistent across different planes within the dIPN (Figure S7). Just as we had observed for the cell bodies, the representation of landmark position by IPN neuropil was not always consistent across fish. While in all fish we detected a consistent retinotopic map (within and across planes), this also appeared phase-shifted by a constant offset in some fish (Figure 4E).

### Representation of landmark position in the habenula

Given that the habenula is the most prominent input to the IPN [47–50], and that the left habenula neurons are known to respond to whole field changes in luminance levels [54–56], we wanted to investigate the left habenula as a potential source for light responses in the IPN.

The visual receptive field properties of left habenula neurons have not been characterized, so we next imaged the habenula while showing the fish the light bar stimuli as before. We found a population of neurons in the habenula that reliably responded to this stimulus (Figure 4F-G). In line with previous studies, a large fraction of left habenula neurons reliably responded to the light stimulus (30 ± 10%, mean ± sd), and a smaller number in the right habenula (10 ± 7%, mean ± sd). In the left habenula, we could detect neurons that had a receptive field localized in the azimuthal direction (Figure 4H, Figure S8). Unlike the responses in the IPN, we could not detect any organization of this azimuthal position representation in the habenula (Figure 4G). It appears that as for visual motion (Figure 1C-D), habenula neurons that respond to light in a particular position are not topographically organized.

### The habenula is not the source of visual motion information to the aHB and IPN

Our data shows that habenular neurons respond to visual motion and that their axons in the IPN show topographically organized tuning to visual motion direction (Figure 1, Figure 3). Given that the habenula projects to both the IPN and aHB, we next wanted to check if it is the left habenula that relays visual motion information to these structures. We used lightsheet microscopy to image a group of fish before and after chemogenetic habenula ablations. In these experiments we used triple transgenic fish; expressing Tg(elavl3:H2B-GCaMP6s), Tg(16715:Gal) and Tg(UAS:Ntr-mCherry). In these fish nitroreductase (Ntr) is expressed exclusively in habenula neurons, such that in the presence of nifurpirinol (NFP) habenula cells will be ablated. As controls, we used siblings of ablated fish that were found not to express Ntr at all (from now on referred to as treatment controls). Figure 5A shows a whole brain stack of a fish brain before and after NFP treatment. Following overnight treatment with NFP, habenula neurons are gone, as well as their axon terminals in the IPN (Figure 5B).

**Figure 5:**
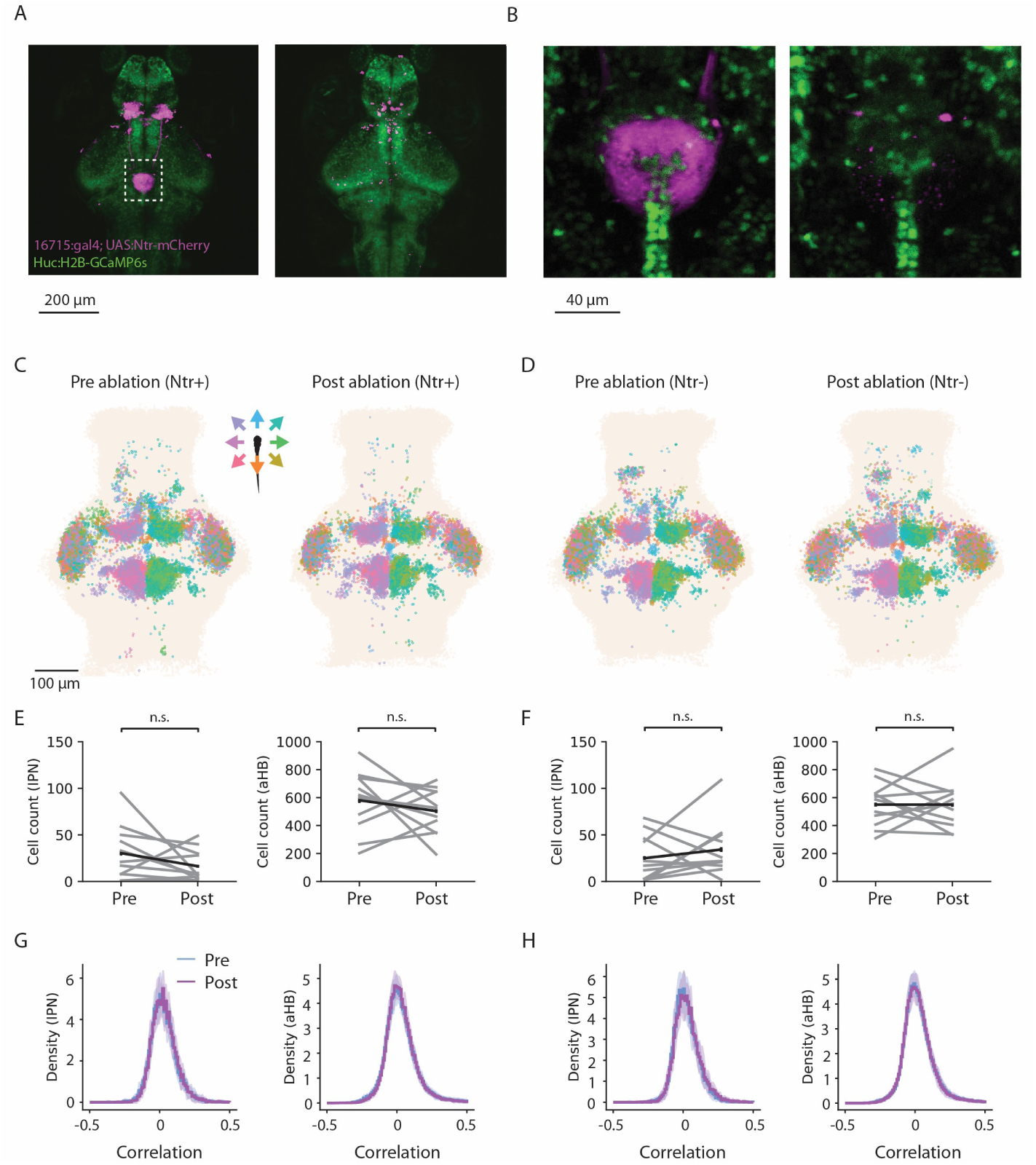
The habenula does not provide visual motion information to the IPN and aHB. (A) Z projection of a confocal stack of a Tg(elavl3:H2B-GCaMP6s; 16715:Gal4; UAS:Ntr-mCherry) fish before (left) and after (right) NFP treatment. (B) Same as A but zoomed in view of the IPN. (C) Pooled ROIs from 11 fish - each neuron is colored according to its correlation value with rightward motion before (left) and after (right) habenula ablation. Only reliably responding neurons are shown (top 5% of reliability index). (D) Same as C but for control fish not expressing Ntr in the habenula (n=11). (E) Number of reliably respondingcellsinthe IPNandaHBbeforeandafterhabenulaablation(distributionsarenotsignificantlydifferent, Wilcoxon signed-rank test, p(IPN)=0.09, p(aHB)=0.23, n=11). Cell count was done by counting all reliably responding neurons in the IPN/aHB (neurons with a reliability threshold > 0.25). (F) Same as E but for control fish not expressing Ntr (distributions are not significantly different, Wilcoxon signed-rank test, p(IPN)=0.84, p(aHB)=0.48, n=11). (G) Distribution of correlation values with all regressors before and after ablation for the IPN and aHB for Ntr+ fish (mean ± sd, n=11). The variances of the distributions before ablation are not significantly larger than those of the distributions post ablation (Wilcoxon signedrank test, p(IPN)=0.25, p(aHB)=0.12). (H) Distribution of correlation values with all regressors before and after ablation for the IPN and aHB for the control group (mean ± sd, n=11). The variances of the distributions before ablation are not significantly larger than those of the distributions post ablation (Wilcoxon signed-rank test, p(IPN)=0.91, p(aHB)=0.28).

Figure 5 shows data from ablated and treatment control fish. Fish from both groups show directional tuning in the tectum, pretectum, aHB, IPN and left habenula before the treatment, as expected. Following NFP treatment, only Ntr+ fish lose visual tuning in the left habenula, indicating that the ablation was successful (Figure 5C). To visualize changes in responses to visual motion we first voxelized the responses in all fish and computed the difference in response amplitude before and after ablation (Figure S9, see Methods). When comparing whole brain response patterns, it appears that habenula ablations do not affect representation of whole field motion in the brain. Specifically, visual tuning in the aHB and IPN remains intact (Figure 5C-F, Figure S10-Figure S12), suggesting that the habenula is not required for representation of visual motion in these regions.

### The habenula provides landmark information to the IPN

Our results show that both habenula and IPN neurons respond to light in localized parts of the visual field (Figure 4). We wanted to check if the habenula is the source of visual landmark information to the dIPN. To do so, we performed genetic ablations of the habenula and recorded activity in the IPN before and after ablation (Figure 6). Before ablation, we could identify IPN neurons that responded to light in a particular position of the visual field in all fish. Following NFP treatment, IPN neurons in Ntr+ fish with ablated habenula showed no response to the presented stimuli (Figure6A-C, FigureS11). IPN neurons in treatment control fish, in which the habenula was left intact, showed similar responses before and after NFP treatment (Figure 6A-C, Figure S13). Combined with our previous results, this data shows that while both the habenula and IPN contain neurons that respond to visual motion and neurons that respond to light position, only the representation in the IPN of the latter requires an intact habenula.

**Figure 6:**
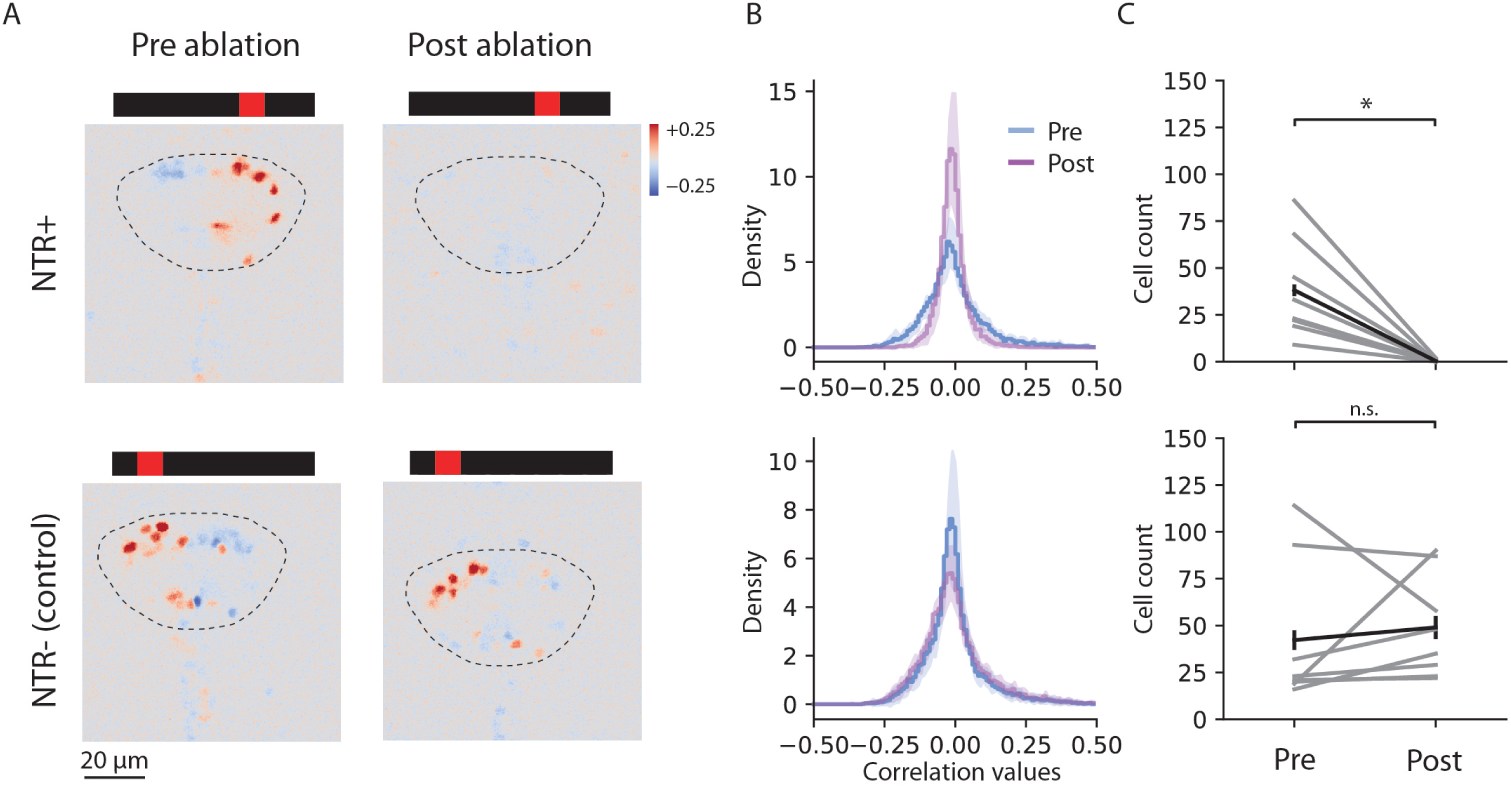
The habenula provides landmark information to the IPN. (A) An example plane showing correlation values of IPN cells with light in a particular position before and after NFP treatment for an Ntr+ (top) and a control (bottom) fish. (B) Top, distribution of correlation values of IPN neurons with the RF regressors before (blue) and after (purple) NFP treatment for Tg(16715:Gal4; UAS:Ntr-mCherry) fish (habenula ablated group, mean ± sd, n=8). The variance of the pre-ablation distribution is significantly largerthan thevarianceofthepostablation distribution (Wilcoxonsigned-rank test, p=0.004). Bottom, distribution of correlation values of IPN neurons with the RF regressors before (blue) and after (purple) NFP treatment for Ntr-fish (control group, mean ± sd, n=8). The variance of the pre-ablation distribution is not significantly larger than the post ablation (Wilcoxon signed-rank test, p=0.9). (C) Number of reliably tuned neurons in the IPN before and after NFP treatment for Ntr+ (top) and a control(bottom) fish. NFP significantly reduces the number of responsive cells in the IPN in the Ntr+ group (Wilcoxon signed-rank test, p=0.004, n=8) but not in the control group (Wilcoxon signed-rank test, p=0.87, n=8). Cell count was done by counting all reliably responding neurons in the IPN (neurons with a reliability threshold > 0.25).

### The habenula is not required for the heading direction network in the aHB

Recently, the habenula was found to directly contact HDNs in the zebrafish aHB [57]. In addition, our data shows that the habenula provides visual information that could be used for landmark based navigation to the IPN (Figure 5). In rodents, a small part of habenular neurons respond to changes in angular heading velocity and could contribute to the construction of the heading signals [41, 47, 58].

We next wanted to check if the habenula is necessary for the heading direction network in the aHB to function in darkness. We used two-photon ablations in order to sever the fasciculus retroflexus (FR), the bundles of axons coming from the habenula to the IPN (Figure 7A). The day after ablation we imaged the fish and checked that the habenular innervation was abolished. We next imaged the aHB of ablated fish and were able to detect the heading direction network (Figure 7B-D). Figure 7C shows the activity of the network in three different fish with ablated habenula. All these fish show an activity bump that moves around the aHB in a way that is correlated with the fish’s estimated heading direction (mean ± sem: -0.88 ± 0.02, n=3), as previously described [6]. This data shows that the habenular input is not necessary for basic functioning of the ring attractor network or for this network to integrate motor commands and estimate the heading direction of the fish. Combined, our results show that while habenula neurons represent different types of signals, their input to the zebrafish heading direction network is specific, in the case of the stimuli we tested, to visual landmark stimuli.

**Figure 7:**
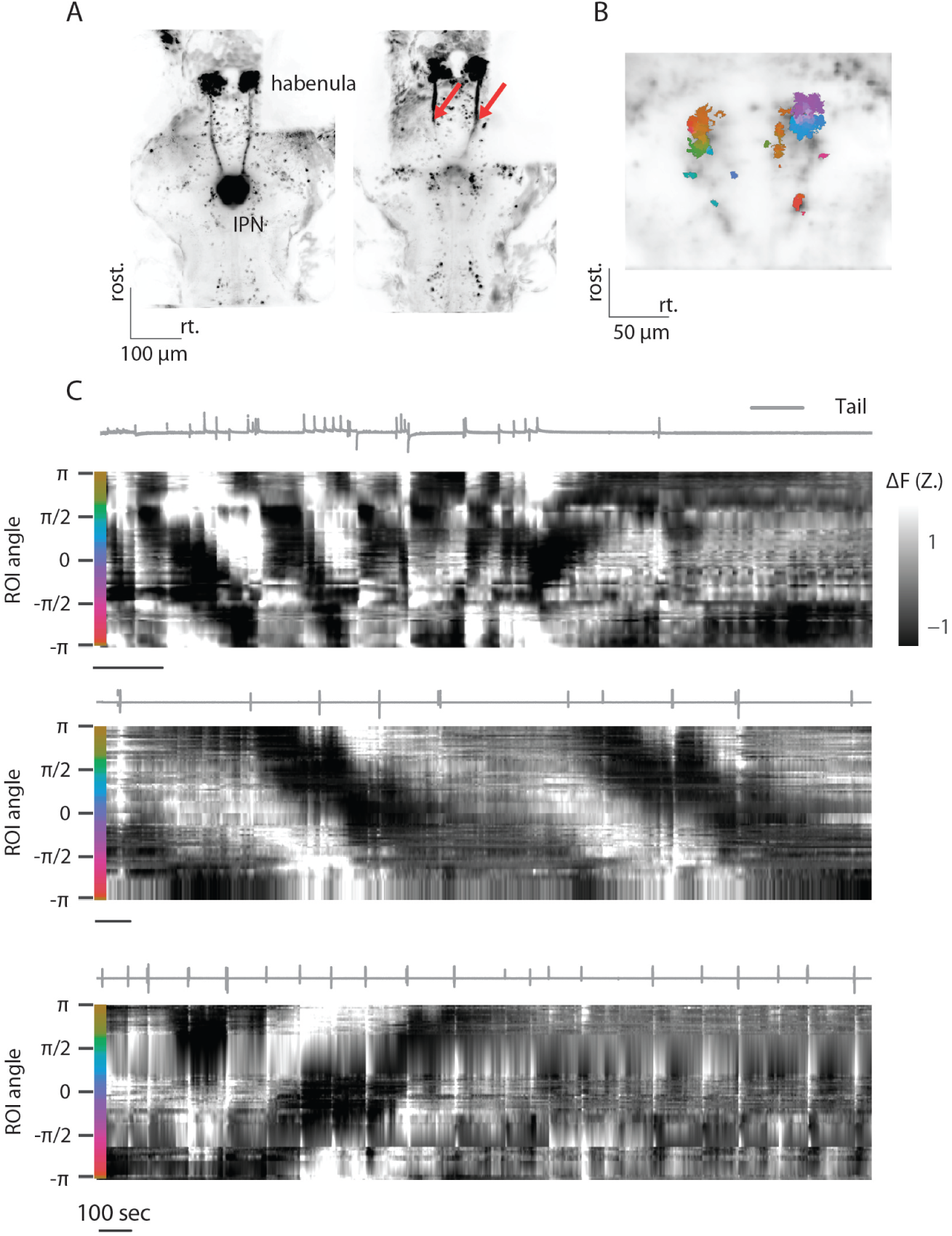
The heading direction network in the aHB functions in the absence of habenular input. (A) Left, z projection of the brain of a Tg(gad1b:Gal4; 16715:Gal4; UAS:GCaMP6s) fish expressing GCaMP6s both in habenular neuron sand GABAergic neurons before two-photon ablation. Right, z projection of the brain of a different fish following a two-photon ablation of the FR. Red arrows indicate the two sites of laser ablation. Note habenular axons are gone from the IPN. (B) aHB anatomy overlaid with HDNs colored according to their phase. (C) Neural dynamics following two-photon ablation of habenular input in three fish. For each fish: top, tail trace showing the motor activity of the fish during the experiment. Bottom, traces of heading direction neurons sorted according to their phase showing the activity bump changing as the fish turns. Black scale bar for each fish indicated 100 seconds of recording.

## Discussion

In this study we investigated the neural representation of visual information in the habenula - IPN - aHB circuit. We focus on two types of visual information that are relevant to navigation: directional whole field motion and landmark position. We find that both signals are represented in the dIPN, the region containing the neuropil of HDNs in zebrafish. These representations are topographically organized in rostro-caudal columns along the dIPN, in a similar manner to the organization of the HD neuropil in this region. We further show that the habenula is necessary for the representation of landmark position in the dIPN, but not for the representation of visual motion or the basic function of the heading direction neurons in the aHB, namely, the integration of motor commands to represent the heading direction of the fish. Our findings suggest that the dIPN is a site of integration of visual information with the heading signal.

The IPN does not contain HDNs, yet it is a core part of the vertebrate heading direction system. In mammals it is bidirectionally connected with the DTN [41] and in zebrafish it is the region in which HDNs are connected to one another [6]. Furthermore, IPN lesions in rats lead to impairment of both landmark based navigation and path integration, indicating the importance of this region in these processes [59]. IPN lesions also affect the stability of HDNs in the thalamus, suggesting that its effects are not restricted to the DTN but encompass the broader HD network [60].

We show that in the aHB, visual motion and heading direction are represented by two separate populations. We further show that representation of visual motion in GABAergic neuropil in the dIPN, most of which comes from the aHB, has a similar structure to the representation of heading direction. In both cases, tuning appears in a columnar structure. Furthermore, the tuning appears in both sides of the dIPN and could be the result of aHB neurons that have a dendrite on one side and an axon on the other. Recent papers have shown different populations of cells in the zebrafish aHB that differ by expression of biochemical markers or morphology [57, 61]. Interestingly, cells that express different markers were found to target different regions of the IPN and show different activity patterns [61]. Future studies are needed to conclude if this is also the case for the HDNs and the visually tuned neurons in this region.

Our data shows that different types of visual information are represented in the dIPN in columns. We show that the anatomical organization of directional visual motion in the dIPN is consistent across fish, unlike the HD signal. When imaging habenular axons in the dIPN or the neuropil of dIPN neurons we see that each direction is represented on a single stripe in the rostro-caudal axis. The topographic organization of tuning in habenular axons is surprising given their morphology. Habenular axons wrap around the IPN and do not form localized projections [53]. Previous studies have shown that the activity of habenula axons in the IPN is shaped by axoaxonic synapses [62]. These synapses could belong to IPN neurons, aHB neurons that arborize in the IPN [57] or a different population that targets the IPN such as the nucleus incertus [61].

The columnar structure in which IPN responses are organized is very reminiscent of the insect central complex. The insect central complex is also a site of integration of different signals that are relevant for navigation, such as heading direction and the animals traveling direction [39, 40]. While the insect HDNs themselves are not sensitive to translational visual motion, another type of cells in this region responds to visual motion in a particular direction [39, 40]. Both types of cells tile regions of the central complex, their neuropil forming columns, each tuned to a different heading/traveling direction. Our data shows that similar organization exists also in the vertebrate brain.

In many animals, the heading direction signal is tethered to the outside world and can be updated by shifting a landmark to a new position [3, 17]. This is crucial for navigation, as while most animals are capable of using path integration for navigation, this mechanism is prone to suffer from accumulating errors. The strong effect of visual landmarks on spatial sensation is evident not only in the heading direction network, but also in heading-direction-modulated place cells in the rodent hippocampus [63]. The directional tuning of place cells is preserved in virtual reality, demonstrating its dependence on visual cues rather than vestibular information [63]. Furthermore, the visual tuning in the hippocampus is also present in immobile rats presented with a moving bar of light [64]. The existence of visual tuning, independent of the rat motor actions, demonstrates that the control of visual landmarks on spatial orientation extends beyond the classic heading direction network [64].

In insects, the tethering of the heading direction signal to the external visual environment is done by ring neurons: neurons that show receptive field responses and carry this information to the ellipsoid body [30, 32]. Unlike many other neurons in the insect heading direction system, ring neurons do not tile a specific wedge of the ellipsoid body, but rather form synapses throughout the structure. Through yet unknown mechanisms of plasticity, these neurons manage to convey information about landmark position to specific HDNs in different environments [30].

The morphology of the larval zebrafish habenula neurons is strikingly similar to the morphology of the insect ring neurons. They both have a single axon that wraps around the target structure and forms synapses in a non-localized manner [38, 53]. Interestingly, habenula ablation in rats impairs several cognitive functions, one of which is landmark based navigation [65]. Here we show that a group of habenula neurons, similarly to the drosophila ring neurons, respond to light in a particular part of the visual field. We further show that ablating the habenula results in absence of landmark responses in the IPN, showing that habenular input is necessary for this representation. These results imply that a group of habenula neurons provide landmark related information to the dIPN, where it is poised to be integrated into the zebrafish heading direction system. The similarities between larval zebrafish habenula neurons and drosophila ring neurons suggests that this information could be integrated via similar mechanisms, revealing further potential analogies between the insect and vertebrate heading direction systems [66].

## Acknowledgements

We thank all the members of the Portugues lab members for their input. H.L. would like to thank Inbal Shainer and Barak Shalom for helpful discussions. This research was funded by the German Research Foundation (DFG) under Germany’s Excellence Strategy within the framework of the Munich Cluster for Systems Neurology (EXC 2145 SyNergy, identifier 390857198) and through the “Enhanced resolution microscopy” project DFG – Projektnummer 518284373, by the Volkswagen Stiftung via a Life? grant and by the Max Planck Foundation.

## Methods

### Zebrafish husbandry

All procedures related to animal handling were conducted following protocols approved by the Technische Universität München and the Regierung von Oberbayern (TVA # 55-2-1-54-2532-101-12 and TVA ROB-55.2-2532.Vet_02-24-5). Adult zebrafish (Danio rerio) from Tüpfel long fin (TL) strain were kept at 27.5-28°C on a 14/10 light cycle, and hosted in a fish facility that provided full recirculation of water with carbon-, bio- and UV filtering and a daily exchange of 12% of water. Water pH was kept at 7.0-7.5 with a 20 g/liter buffer and conductivity maintained at 750-800 µS using 100g/liter. Fish were hosted in 3.5 liter tanks in groups of 10 to 17 animals. Adults were fed with Gemma micron 300 (Skretting) and live food (Artemia salina) twice per day and the larvae were fed with Sera micron Nature (Sera) and ST-1 (Aquaschwarz) three times a day.

All experiments were conducted on 5-10 dpf larvae of yet undetermined sex. The week before the experiment, one male and one female or three male and three female animals were left breeding overnight in a breeding tank (Tecniplast). The day after, eggs were collected in the morning, rinsed with water from the facility water system, and then kept in groups of 20-40 in 90 cm Petri dishes filled with 0.3x Danieau’s solution (17.4 mM NaCl, 0.21 mM KCl, 0.12 mM MgSO4, 0.18 mM Ca(NO3)2, 1.5 mM HEPES, reagents from Sigma-Aldrich) until hatching and in groups of 20 larvae in water from the fish facility afterwards. Larvae were kept in an incubator that maintained temperature at 28.5°C and a 14/10 hour light/dark cycle, and their solution was changed daily. At 4 or 5 dpf, animals were lightly anesthetized with Tricaine mesylate (Sigma-Aldrich) and screened for fluorescence under an epifluorescence microscope. Animals positive for GCaMP6s/ mCherry fluorescence were selected for the imaging experiments. Animals older than 5 dpf were kept in small breeding tanks and fed daily with Sera micron Nature and Rotifers.

### Transgenic animals

Tg(elavl3:H2B-GCaMP6s) fish were used for whole brain imaging experiments [67]. Imaging of GABAergic neurons was done using double transgenic animals expressing Tg(gad1b/GAD67:Gal4-VP16)mpn155 (referred to as Tg(gad1b:Gal4)) which drives expression in a subpopulation of GABAergic cells under gad1b regulatory elements [68] and Tg(UAS:GCaMP6s)mpn101 [69]. For genetic ablation experiments, we used a transgenic line expressing three different elements: Tg(elavl3:H2B-GCaMP6s), Tg(16715:Gal4) and Tg(UAS:Ntr-mCherry) [70]. For imaging IPN neurons and their neuropil we used the double transgenic animals expressing Tg(S1168t:Gal4) [71] and Tg(UAS:GCaMP6s)mpn101. For imaging the pan-neuronal neuropil in the IPN we imaged Tg(elavl3:GCaMP6s) fish [72]. All the transgenic animals were also mitfa-/- and thus lacked melanophores [73].

### Lightsheet imaging

Lightsheet experiments were done as previously described [6, 74]. Briefly, animals were embedded in 2-2.5% low-melting point agarose (Thermofisher) in a custom lightsheet chamber with a glass coverslip sealed on the sides in the position where the beams of the lightsheet enters the chamber, and a square of transparent acrylic on the bottom, for behavioral tracking. The chamber was filled with water from the fish facility system and agarose was removed along the optic path of the lateral laser beam (to prevent scattering), and around the tail of the animal, to enable movements of the tail. After embedding, fish were left recovering 1 to 6 hours before the imaging session.

Imaging experiments were performed using a custom-built lightsheet microscope [75] as previously described [6, 74]. In whole brain experiments, 20 planes were acquired over a range of approximately 250 µm, slightly adjusted for every fish. The resulting imaging data had a resolution of 10 × 0.6 × 0.6 µm/voxel, and a temporal resolution of 2 Hz. In experiments imaging only the aHB, 8 planes were acquired over a range of approximately 100 µm, slightly adjusted for every fish. The resulting imaging data had a resolution of 10 × 0.6 × 0.6 µm/voxel, and a temporal resolution of 5 Hz.

### Two-photon microscopy

Two-photon experiments were done as previously described [6, 45]. Briefly, animals were embedded in 2-2.5% low-melting point agarose (Thermofisher) in 30 mm petri dishes. The agarose around the tail, caudal to the pectoral fins, was cut away with a fine scalpel to allow for tail movement. The dish was placed onto an acrylic support with a light-diffusing screen and imaged on a custom-built two-photon microscope. In experiments in which visual stimuli were projected in the front visual field of the fish, the front half of the petri dish was covered with a light diffusing paper. The custom Python package brunoise was used to control the microscope hardware [76].

Full frames were acquired at 3 Hz in four, 0.83 μm-spaced interlaced scans, which resulted in x and y pixel dimensions of 0.3 - 0.6 μm (varying resolutions depending on field of view covered). After acquisition from one plane was done, the objective was moved downward by 2 - 8 μm and the process was repeated.

Visual stimuli were generated using a custom written Python script with the Stytra package [77], and were projected at 60 frames per second using an Asus P2E microprojector and a red long-pass filter (Kodak Wratten No.25) to allow for simultaneous imaging and visual stimulation. Fish were illuminated using infrared light-emitting diodes (850 nm wavelength) and imaged from below at up to 200 frames per second using an infrared-sensitive charge-coupled device camera (Pike F032B, Allied Vision Technologies). Tail movements were tracked online using Stytra.

### Tail tracking and stimulus presentation

To monitor tail movements during the imaging session, an infrared LED source (RS Components, UK) was used to illuminate the larvae from above. A camera (Ximea, Germany) with a macro objective (Navitar, USA) was aimed at the animal through the transparent bottom of the lightsheet chamber with the help of a mirror placed at 45° below the imaging stage. A longpass filter (Thorlabs, USA) was placed in front of the camera. A projector (Optoma, Taiwan) was used to display visual stimuli; light from the projector was conveyed to the stage through a cold mirror that reflected the projected image on the 45°-mirror placed below the stage. The stimuli were projected on a white paper screen positioned below the fish, with a triangular hole that kept the fish visible from the camera. The behavior tracking part of the rig was very similar to the setup for restrained fish tracking described in [77].

Frames from the behavioral camera were acquired at 400 Hz and tail movements were tracked online using Stytra [77] with Stytra’s default algorithm to fit to the tail 9 linear segments. The “tail angle” quantity used for controlling the closed-loop was computed online during the experiment in the Stytra program as the difference between the average angle of the first two and last two segments of the tail and saved with the rest of the log from Stytra. The stimulus presentation and the behavior tracking were synchronized with the imaging acquisition with a ZMQ-based trigger signal supported natively by Stytra.

### Visual stimulation

To study neural responses to visual motion we presented fish with a pink noise pattern from below. The pattern could move in 8 different directions with even 45 degrees spacing. In each trial, the pattern moved in one direction (chosen randomly) for 10 seconds and then paused for 5/10 seconds before the next trial.

To study neural responses to landmark position, we presented fish with a bar of light in its front visual field. The fish was embedded in a plastic dish and the front half of the dish was covered with filter paper. A projector was placed in front of the dish as illustrated in Figure 5. A red rectangle appeared in one of 8 possible locations in the fish’s front visual field in a random order. The red rectangle appeared for 5/10 seconds and then disappeared for 5/10 seconds before appearing again in a new location. For these experiments we chose to use two-photon microscopy, as in a lightsheet microscope the fish can see the blue laser which could be perceived as an additional landmark.

### Chemical ablations

For chemical ablations of habenular neurons we used fish expressing three transgenic elements: Tg(elavl3:H2B-GCaMP6s), Tg(16715:Gal4), and Tg(UAS:Ntr-mCherry). In these fish Ntr is only expressed in the habenula. Ntr+ and Ntr-fish were imaged before ablation at 5-6 dpf. Following the first imaging session, fish were left in a 5 µM NFP solution in a light protected box for 12-14 hours as previously described [70, 78]. Fish were washed several times and left to recover for 24 hours before they were imaged again at 7-8 dpf.

### Confocal experiments

For confocal experiments, larvae were embedded in 2% agarose and anesthetized with Tricaine mesylate (Sigma-Aldrich). Whole brain stacks of 5 dpf fish expressing Tg(elavl3:H2B-GCaMP6s), Tg(16715:Gal4), and Tg(UAS:Ntr-mCherry) transgenes were acquired using a 10x water immersion objective (NA = 0.45) with a voxel resolution of 1 x 0.83 x 0.83 (LSM 880, Carl Zeiss, Germany). Stacks of the IPN were acquired using a 20x water immersion objective (NA = 1) with a voxel resolution of 1 x 0.28 x 0.28. The fish were freed from the agarose, treated with NFP for 16 hours and imaged again with identical parameters on 7 dpf.

### Two photon laser ablations

For two-photon laser ablations of the fasciculus retroflexus (FR) tracts of habenular axons going in the IPN we used fish expressing three transgenic elements: Tg(gad1b:Gal4), Tg(16715:Gal4), and Tg(UAS:GCaMP6s). In these fish GCaMP6s is expressed in the habenula and in GABAergic neurons in the brain. The labeling of the FR tracts allowed us to target the two-photon laser to a thin section (5x1 µm) of each of the two tracts. This section was targeted with the two-photon laser at 130 mW for 200 ms. Each tract was cut twice, about 20 and 40 µm rostral to the IPN. Following laser ablation, the fish were freed from agarose and left to recover overnight at 28 degrees with available food. The following day, fish health was assessed and only fish which were active and fed were chosen for the following experiments. Fish were embedded again and a whole brain stack was acquired to ensure that habenular axons were ablated and missing from the IPN. Next the GABAergic neurons in the aHB were imaged for detection of the heading direction network in ablated fish.

### Imaging data preprocessing

The imaging stacks were saved in hdf5 files and then directly fed into suite2p, a Python package for calcium imaging data registration and ROI extraction [79]. We did not use suite2p algorithms for spike deconvolution. Parameters used for registration and source extraction in suite2p can be found in the shared analysis code. The parameters used by the suite2p algorithm were different based on the microscope used and the transgenic line. From the raw F traces saved from suite2p (F.npy file), ΔF/F_baseline was calculated taking F_baseline as the average fluorescence in a rolling window of 900s, to compensate for some small amount of bleaching that was observed in some acquisition. The signal then was smoothed with a median filter from scipy (medfilt from scipy.signal), and Z-scored so that all traces were centered on 0 and normalized to a standard deviation of 1. The coordinate of each ROI was taken as the centroid of its voxels.

### Reliability index

Reliability index was calculated as previously described [80]. Briefly, we calculated for each ROI the average correlation of the responses across all individual presentations of the presented stimuli. To use an objective criterion to select responsive cells, we used Otsu’s method from the SciPy package to set a threshold on the obtained histogram.

### Regressor analysis

Regressor analysis was done as previously described [81]. Briefly, regressors were generated from stimulus related variables. In the whole field visual motion experiments, we constructed 8 regressors, one for each direction of motion. In the landmark experiments we constructed 8 regressors, one for each landmark position. The regressors were convolved with an exponential decay kernel that was found to fit our data. The constructed regressors were correlated with traces extracted from segmented ROIs or individual pixels (depending on the analyzed data).

### Tuning maps

Tuning maps were generated using the results from the regressor analysis described above. For each analyzed pixel/ ROI we had 8 different correlation values with the 8 regressors. For generation of tuning maps, each pixel/ ROI was assigned two values: angle and amplitude. The angle represents the preferred direction of each pixel by indicating which direction elicits the strongest response, and is indicated in the maps by the pixel’s hue. The amplitude represents the magnitude of directional tuning and is indicated in the maps by the pixel’s saturation.

### Quantification of neural tuning to landmark position

To characterize the representation of landmark position in the habenula, tuning curves were generated for each left habenula neuron based on the fluorescence measured when the landmark was presented in different positions in the range ±60 degrees. Each tuning curve was fit with a von Mises function using least squares optimization.

### Voxel-wise differences

For the voxel-wise quantification of neuronal tuning differences as a result of habenula ablations (Figure S9), the MapZeBrain [82] reference brain was first split in cubic voxels of 5 μm per side. Then for each one of the four imaging sessions (Ntr+ and control fish, for both pre- and post-ablation imaging sessions) all detected ROIs were assigned to their corresponding voxel containing them, and the final tuning amplitude for each voxel was computed as the average amplitude from all ROIs contained in it (top and middle rows in Figure S9). The difference in tuning amplitude between pre- and post-ablation imaging sessions were computed as the voxel-wise difference of this value across the two experimental sessions (bottom rows in Figure S9).

### Anatomical registrations

For whole brain lightsheet experiments, all individual brains were registered to the MapZeBrain atlas reference brain [82]. Brain registration was performed on the anatomical stacks obtained from averaging selected frames in the corresponding dataset along the temporal dimension. First, a manual registration was performed using a custom napari-based GUI [83]. Results from this initial alignment were then used as the initial registration and fed into an ANTsPy registration pipeline that included both affine and diffeomorphic transformations [84]. In order to manipulate the different anatomical spaces, the brainglobe-space package from Brain-Globe [85] was used. Coordinates for each ROI were computed as the centroid of all of its encompassing voxels. Registration of Two-photon was done using a similar pipeline. Two-photon datasets were not registered to the MapZeBrain atlas but to one of the fish from the specific dataset.

### Data analysis and statistics

All parts of the data analysis were performed using Python 3.7, and Python libraries for scientific computing, in particular Numpy, Scipy and Scikit-learn. The Python environment required to replicate the analysis in the paper can be found in the paper code repository. All figures were produced using Matplotlib. All the analysis code will be available upon publication.

---

**Figure S1:**
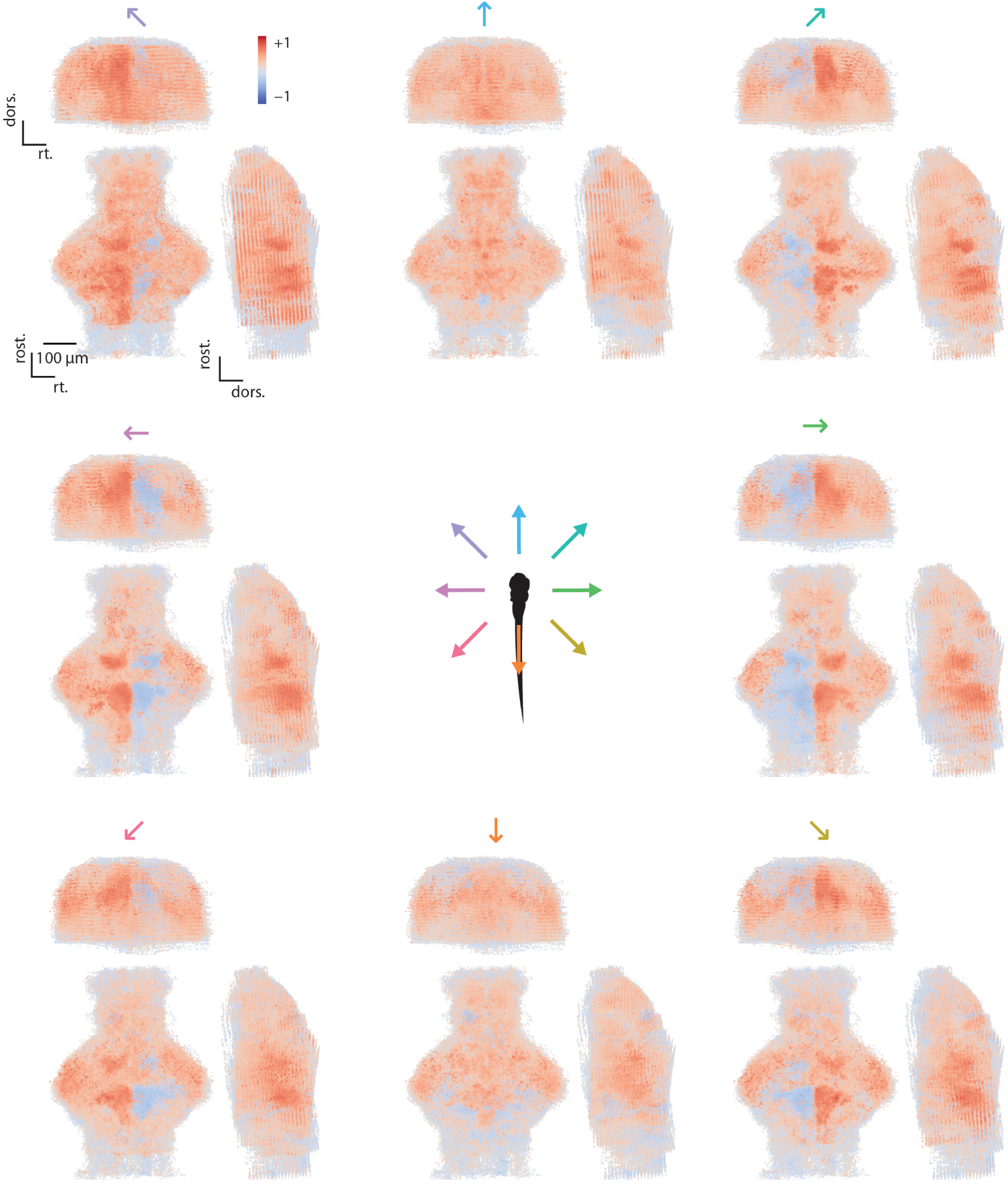
Correlation with visual motion. Correlation maps for all 8 regressors for all fish in the dataset morphed to a reference brain (n=15). In each panel there are three views of the brain (max projections), each neuron is colored according to its correlation value with a particular direction of motion, indicated by the arrow above each plot.

**Figure S2:**
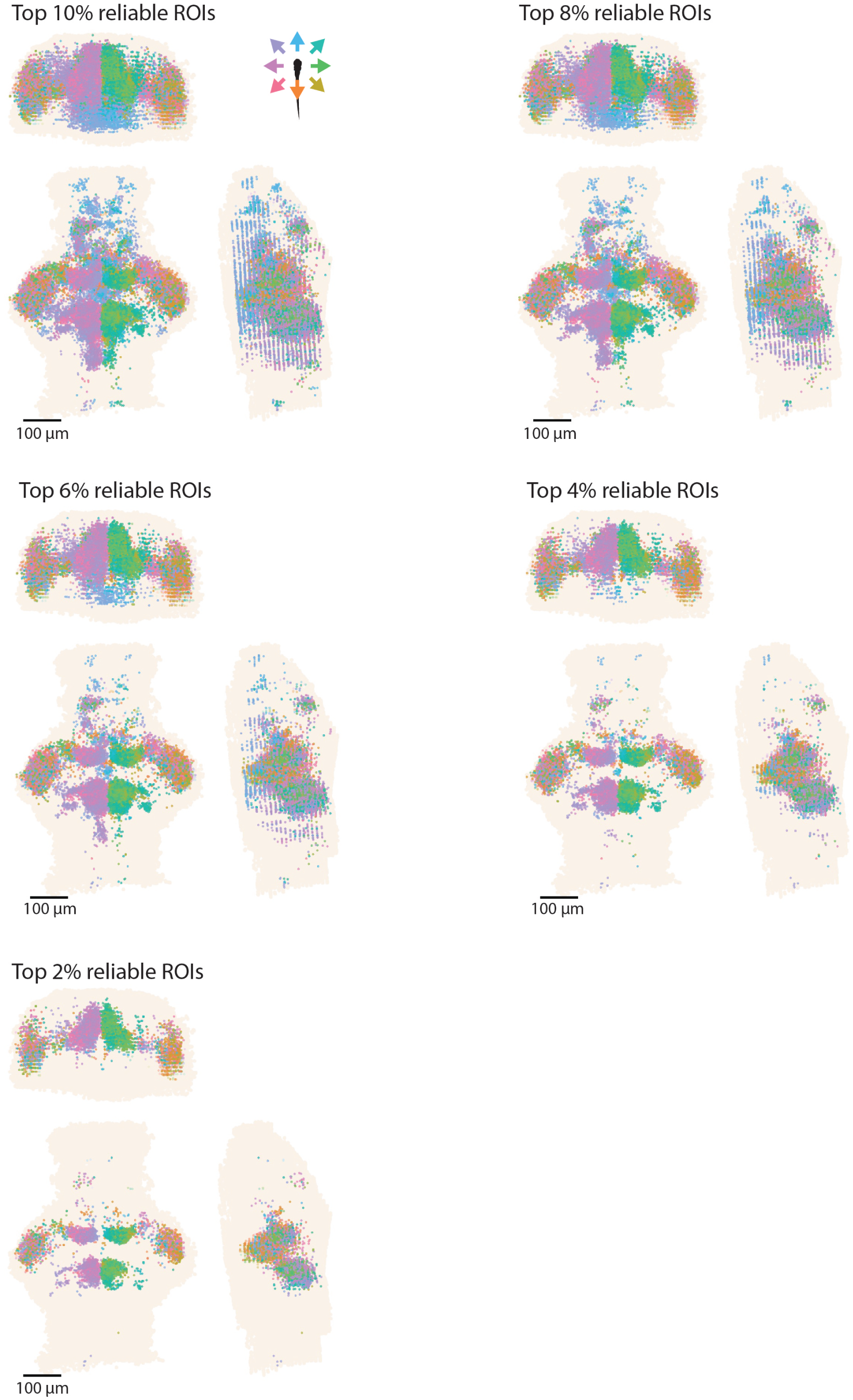
Reliably visually tuned neurons. Tuning maps of all fish in the dataset (n=15), only reliably responding neurons are shown. Each neuron is colored according to the direction it is tuned to. In each plot a different fraction of reliably responding neurons are shown.

**Figure S3:**
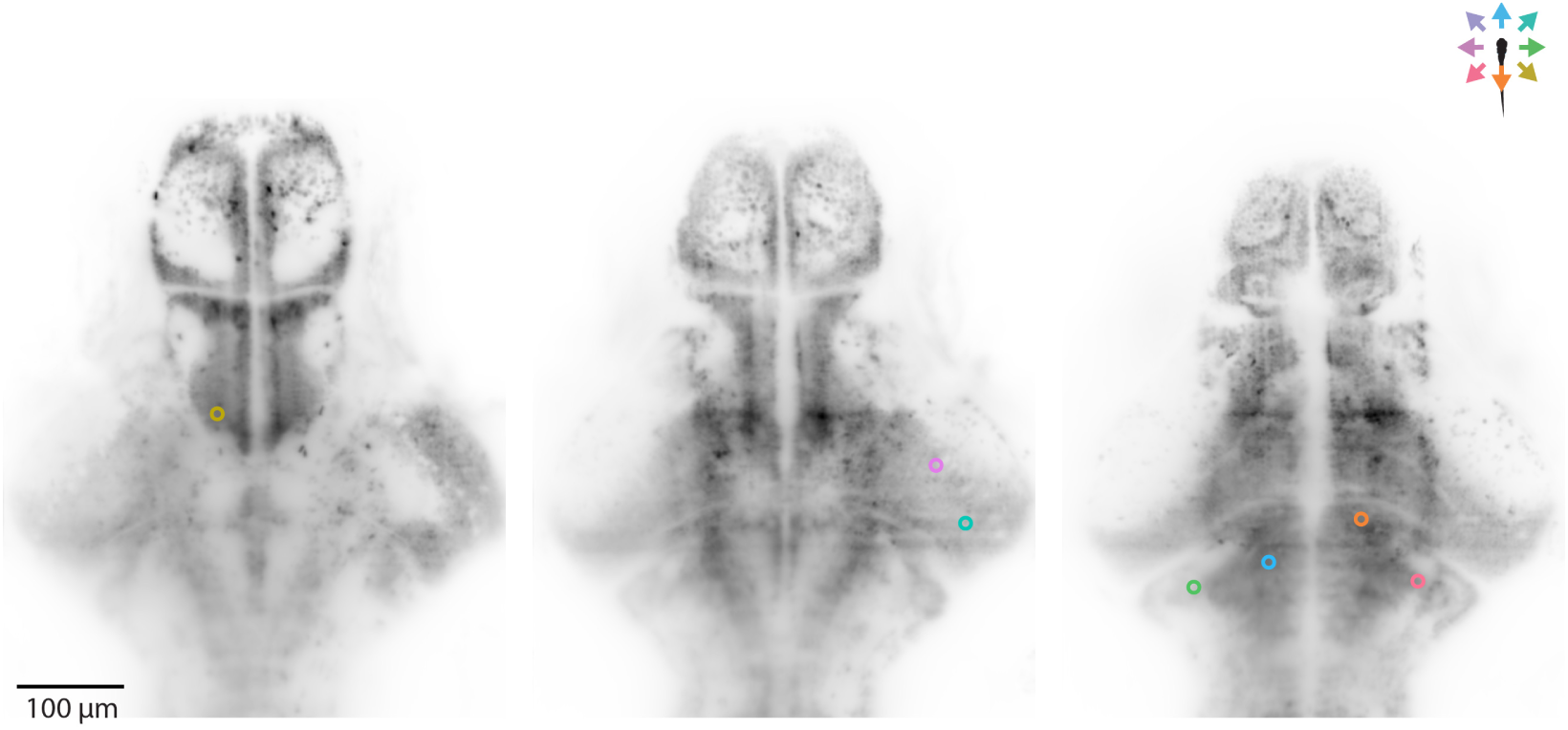
Location of the example neurons shown in Figure 1. Each neuron is colored according to the direction it is tuned to, and its position is plotted over imposed to the anatomical slice containing it (28 μm, z projection of 3 imaging planes).

**Figure S4:**
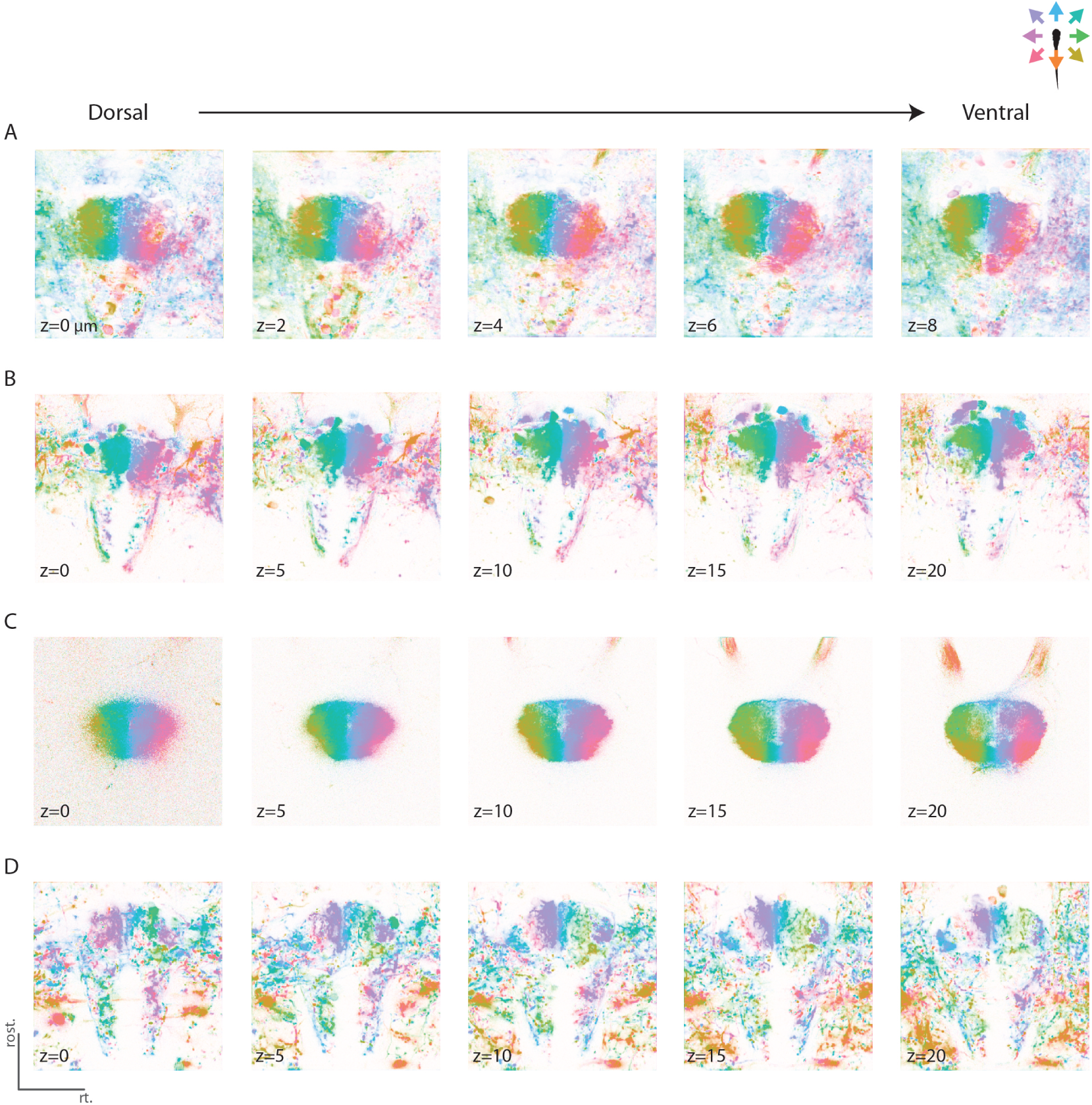
Tuning maps of dIPN neuropil to directional whole field visual motion are consistent across different planes. Example tuning maps from the following lines Tg(elavl3:GCaMP6s) (A), Tg(s1168t:Gal4;UAS:GCaMP6s) (B), Tg(16715:Gal4;UAS:GCaMP6s) (C) and Tg(gad1b:Gal4;UAS:GCaMP6s) (D). Each plane was recorded separately. The z values indicate the distance between different planes (in μm).

**Figure S5:**
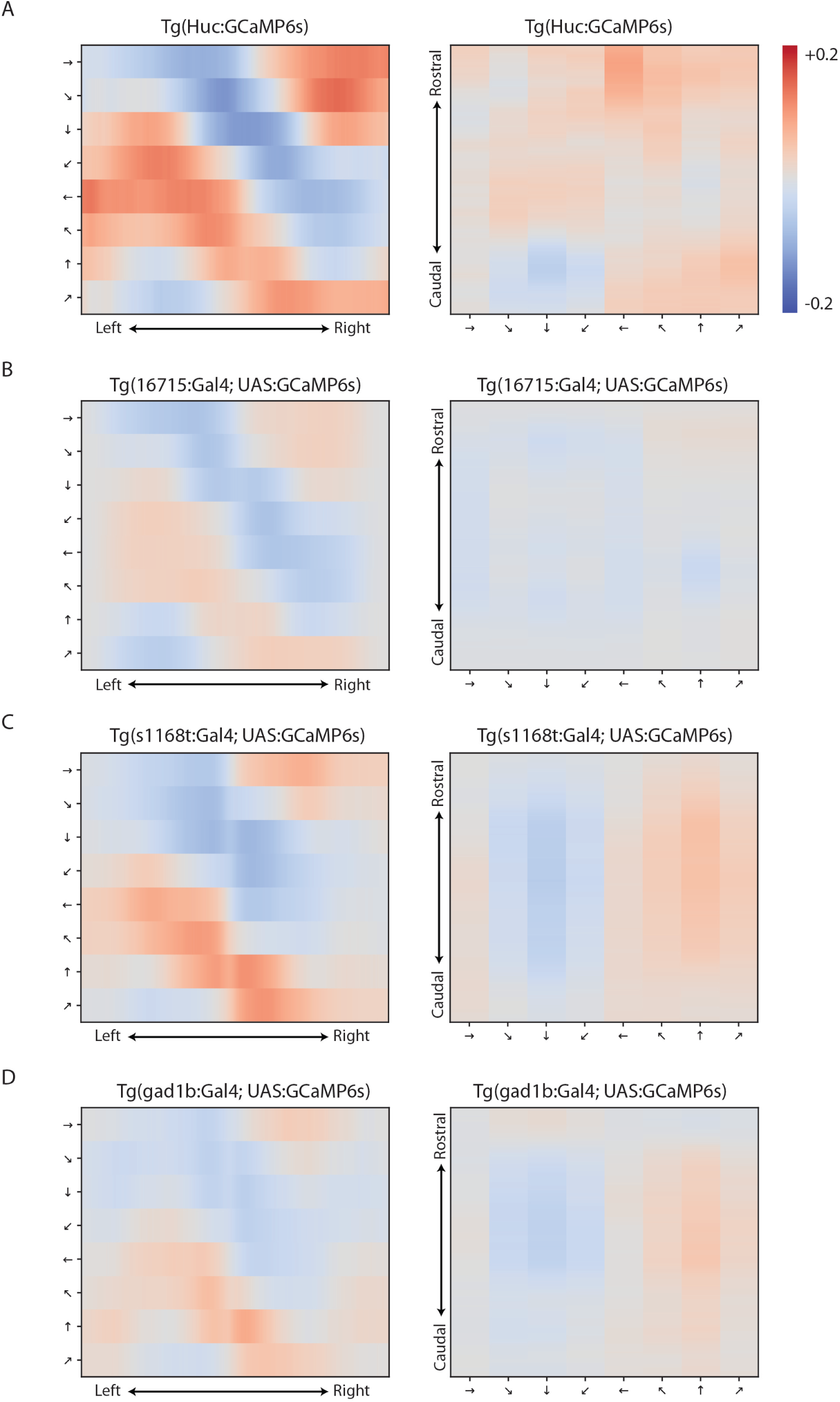
Topographic organization of visual motion changes along the lateral-medial axis in the dIPN. Correlation matrix showing the change in correlation averaged over all pixels in each column along the lateral-medial axis (left) and in each row along the rostro-caudal axis (right) of the dIPN. The correlation matrices were calculated for Tg(elavl3:GCaMP6s) fish (n=4, A), Tg(16715:Gal4; UAS:GCaMP6s) fish (n=12, B), Tg(s1168t:Gal4; UAS:GCaMP6s) fish (n=15, C) and Tg(gad1b:Gal4; UAS:GCaMP6s) fish (n=11, D).

**Figure S6:**
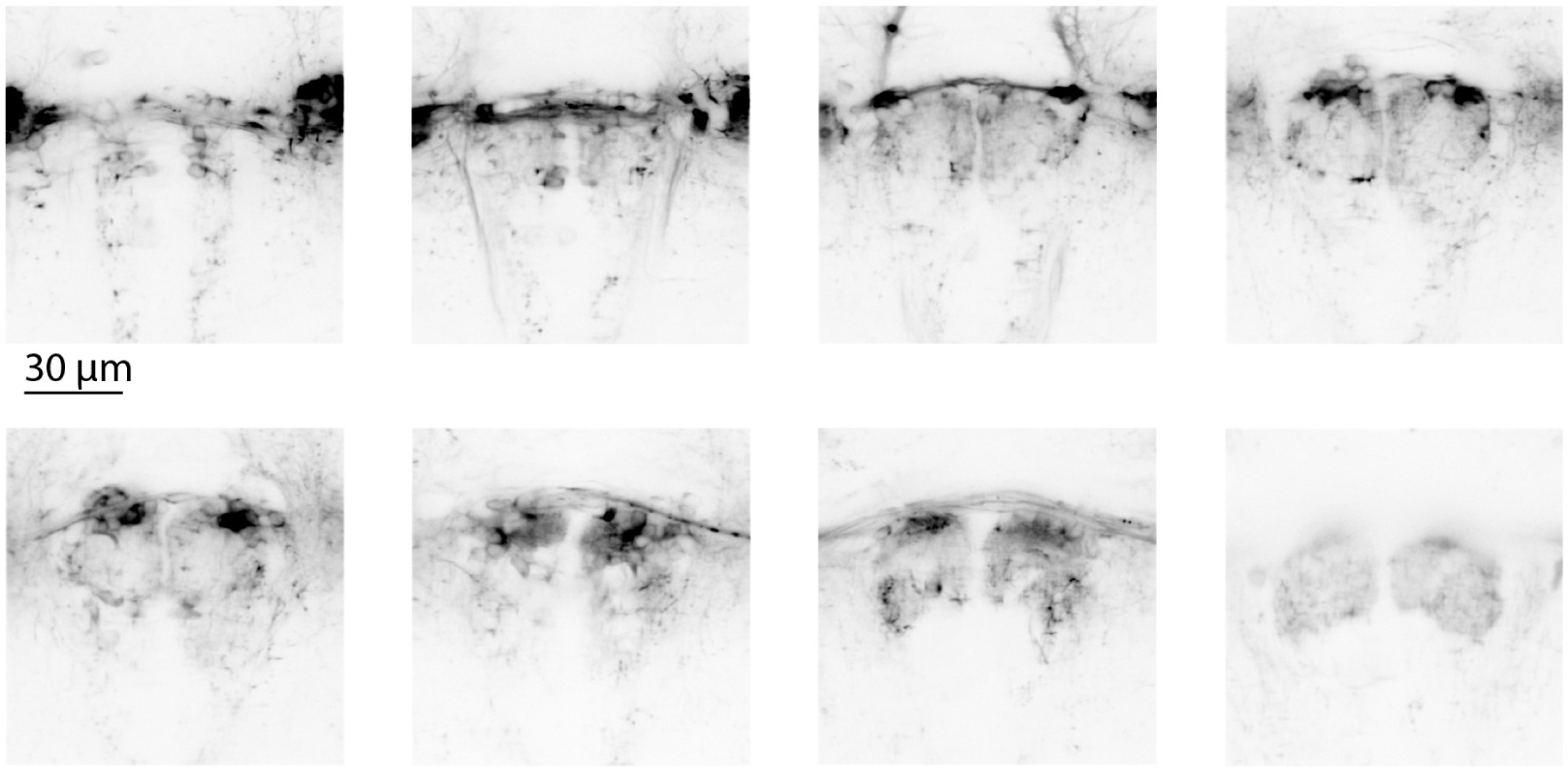
Anatomy of the IPN in the Tg(s1168t:Gal4; UAS:GCaMP6s) line. In addition to expression in eurydendroid cells in the cerebellum and in the tectum [71], this line induces expression of GCaMP6s in a group of IPN cells. The images show the expression pattern in the IPN in different planes, separated by 10 μm from dorsal (top-left) to ventral (bottom-right).

**Figure S7:**
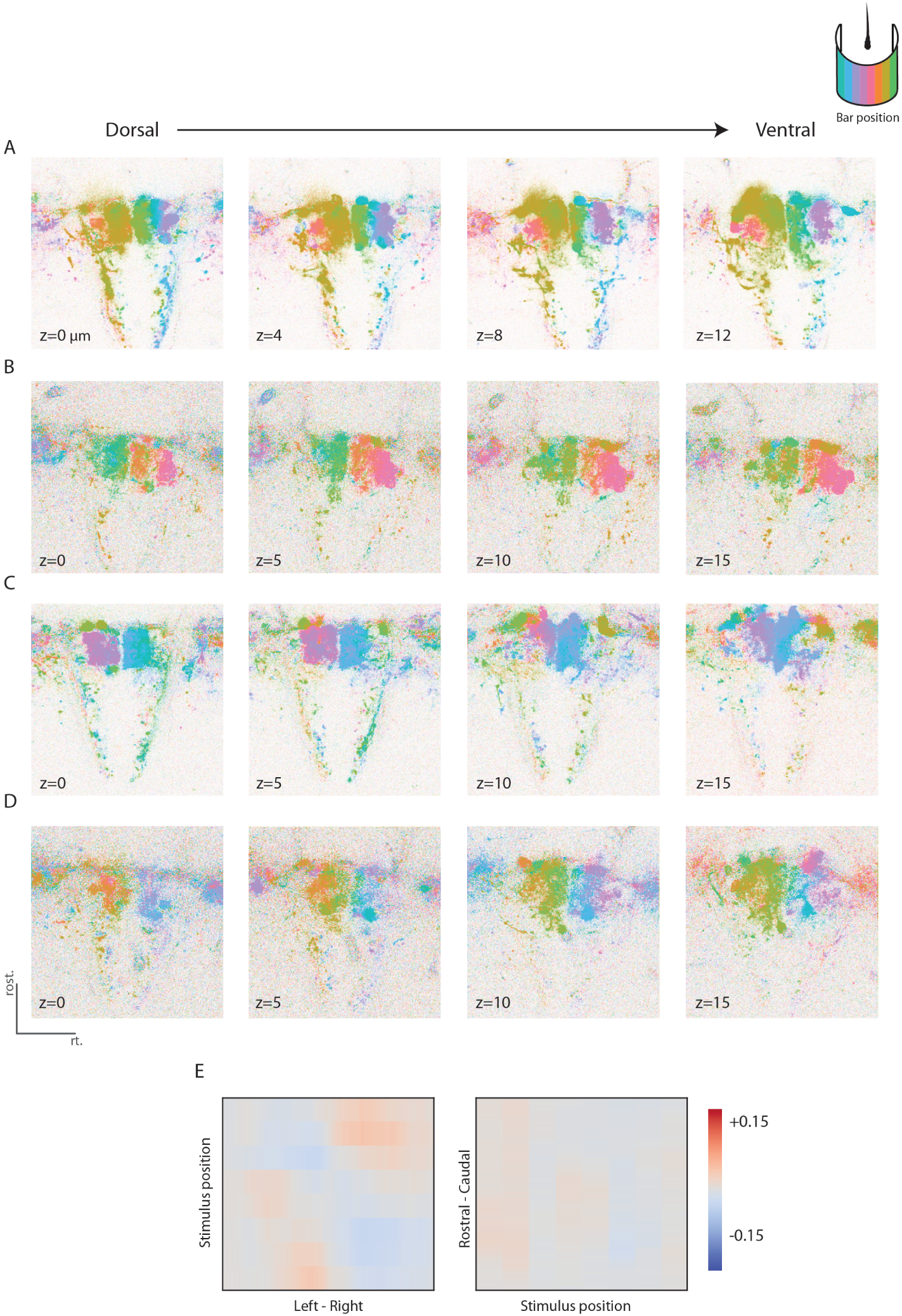
Tuning maps of dIPN neuropil to landmark position are consistent across different planes. (A-D) each row is a different fish (all from the Tg(s1168t:Gal4;UAS:GCaMP6s) fish line). Each plane was recorded separately. The z values indicate the distance between different planes (in μm). E, Correlation matrix showing the change in correlation averaged over all pixels in each column along the lateral-medial axis (left) and in each row along the rostro-caudal axis (right) of the dIPN (n=11).

**Figure S8:**
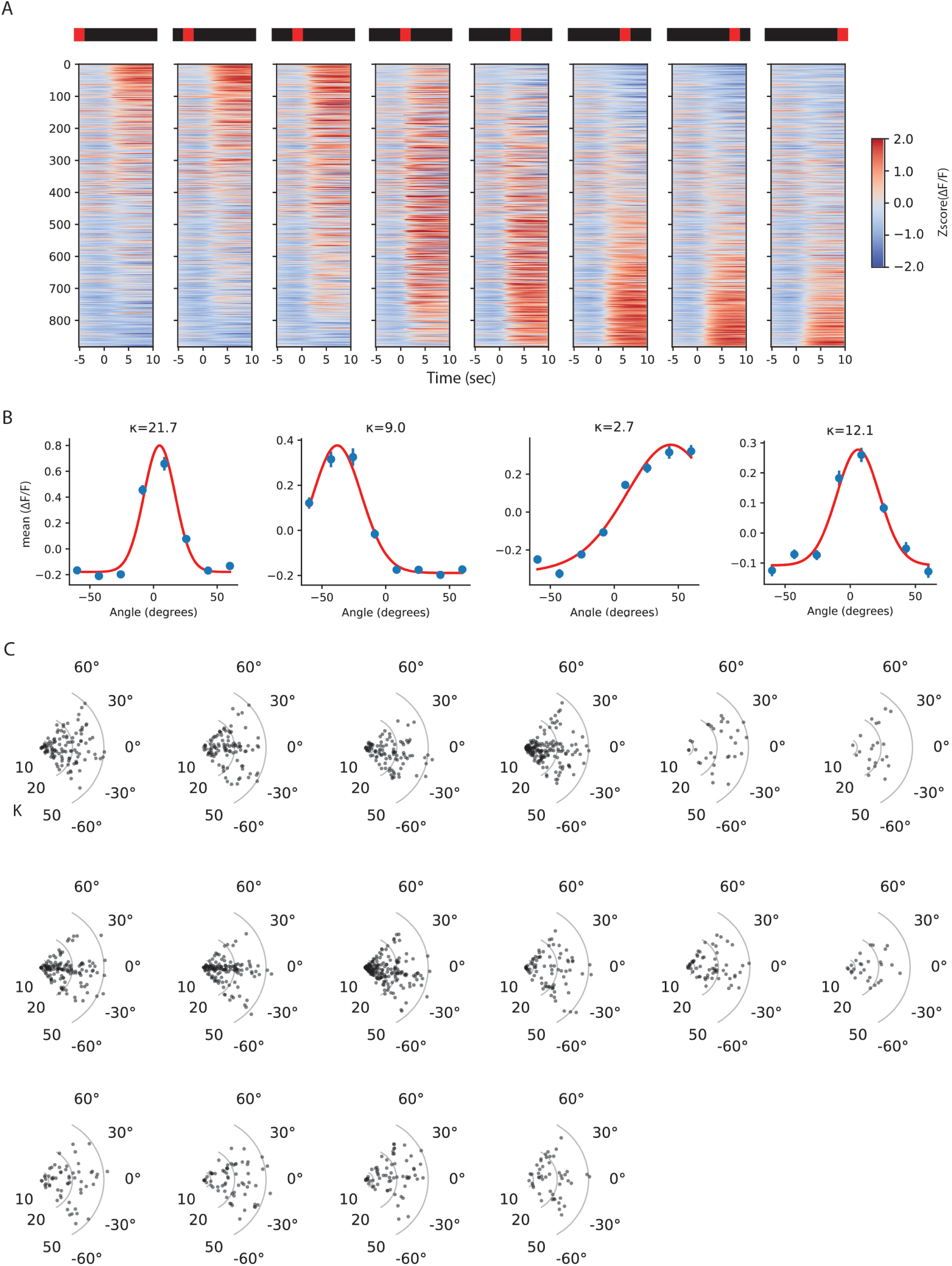
Neurons in the left habenula respond to light bar presented in localized parts of the visual field. (A) Average traces of all reliably responding neurons in the left habenula. Fish were presented with a light bar appearing in different azimuth angles of their visual field. Neurons were sorted according to their center of mass. (B) Tuning curves of 4 example neurons in the left habenula. Each tuning curve was fit with a von Mises function (red line). Kappa values (k), indicating tuning width, are indicated on the individual plot. (C) Distribution of kappa values as a function of the neurons preferred direction, for all left habenula neurons in every fish (n=16 fish). Kappa values are plotted on a logarithmic scale to better visualize the range of tuning widths Fraction of kappa values > 10 in each dataset is 0.56 ± 0.1 (mean ± sd).

**Figure S9:**
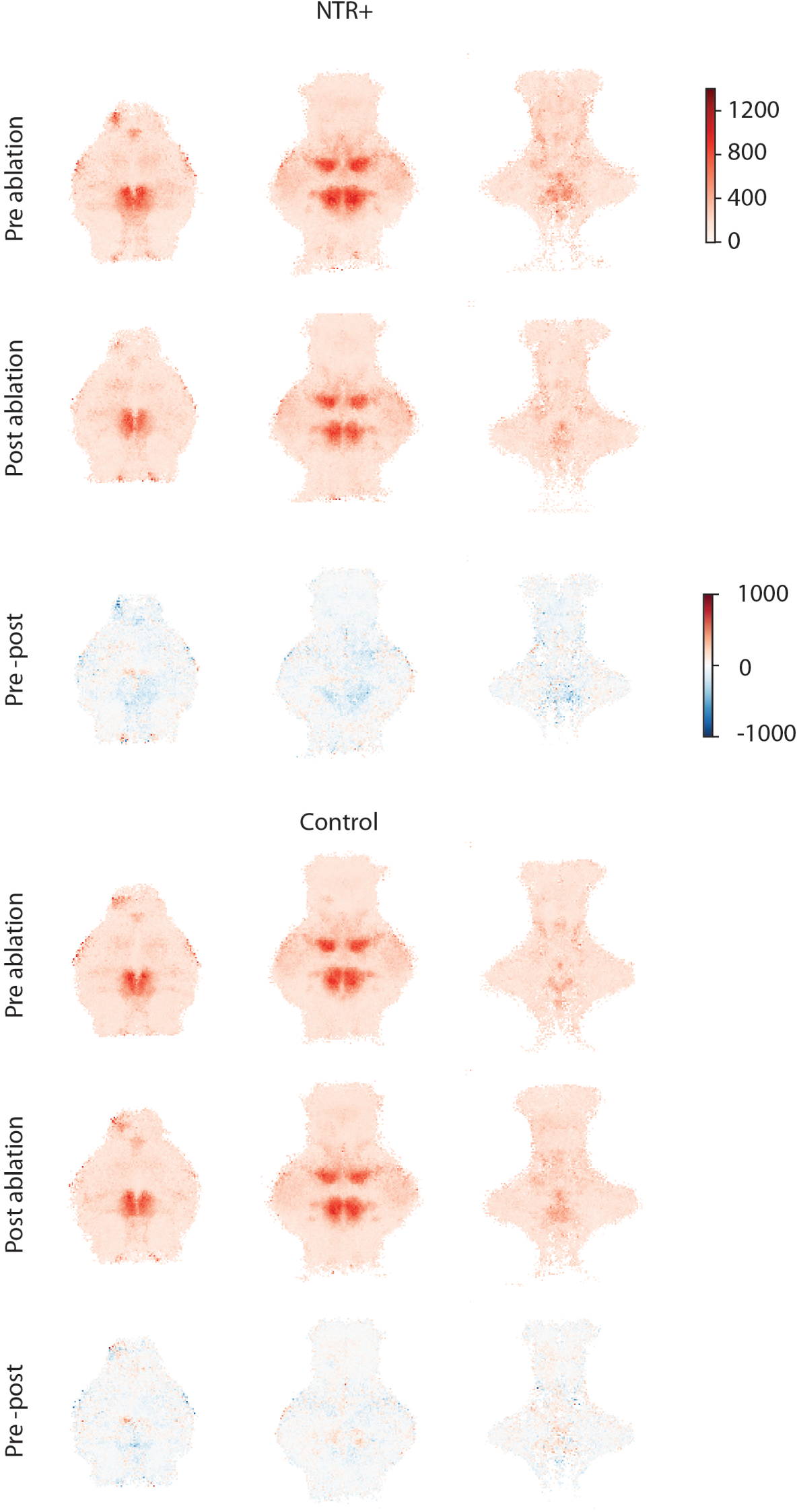
Voxel-wise quantification of tuning amplitude difference as a result of habenular ablation. For both Ntr+ (top panels) and control (bottom panels) fish, the average tuning amplitude of 5 μm cubic voxels is shown during the pre- and post-ablation imaging sessions(top and middle rows respectively). The voxel-wise difference in tuning amplitude across these two sessions is shown in the bottom panels for each experimental condition (see methods). Within each row, the three panels correspond to three anatomical slices of around 120 μm, moving ventrally from left to right.

**Figure S10:**
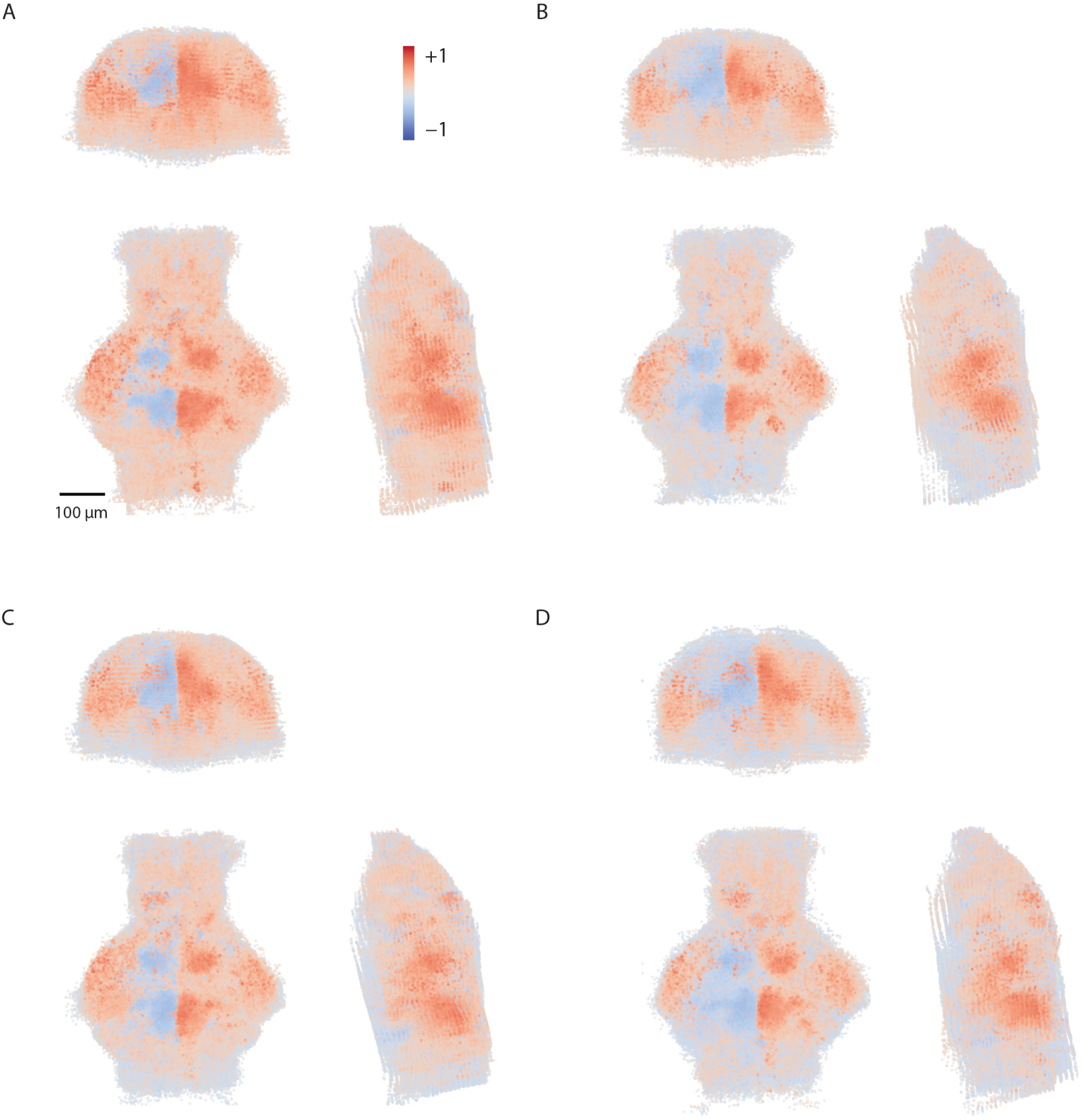
Correlation with rightward motion for all fish before and after habenula ablation. (A) Correlation map for Ntr+ fish before NFP treatment (n=11). (B) Correlation map for the same Ntr+ fish after NFP treatment. (C) correlation map for control fish before NFP treatment (n=11). (D) correlation map for the same control fish after NFP treatment.

**Figure S11:**
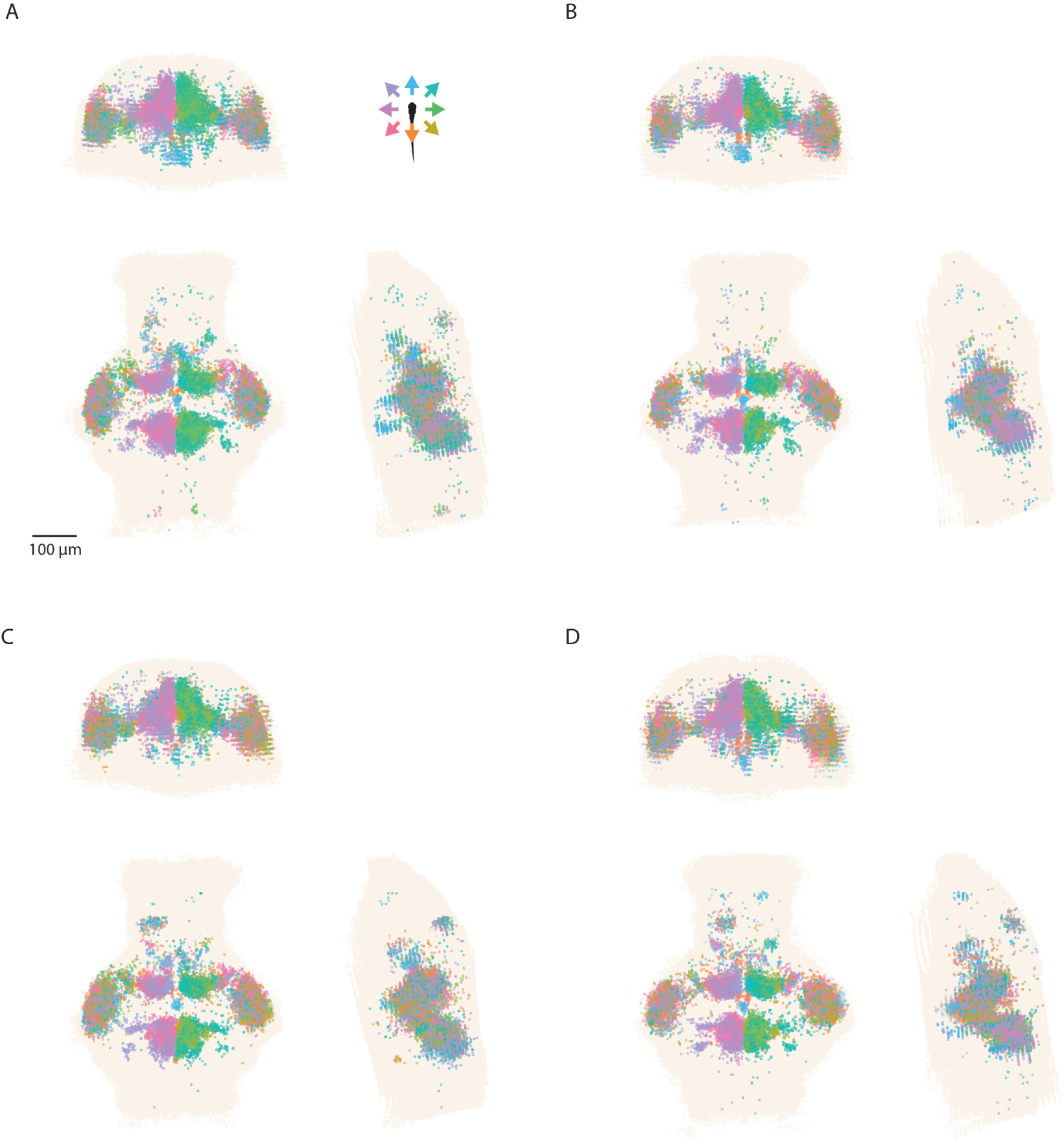
Three views of tuning maps for Ntr+ and control fish before and after habenula ablation. (A) Tuning map of Ntr+ fish before NFP treatment (n=11). Neurons are colored according to their preferred direction, only reliably responding neurons are shown. (B) Tuning map of the same Ntr+ fish after NFP treatment. (C) Tuning map of control fish before NFP treatment (n=11). (D) Tuning map of the same control fish after NFP treatment. Neurons are colored according to their preferred direction, only reliably responding neurons are shown.

**Figure S12:**
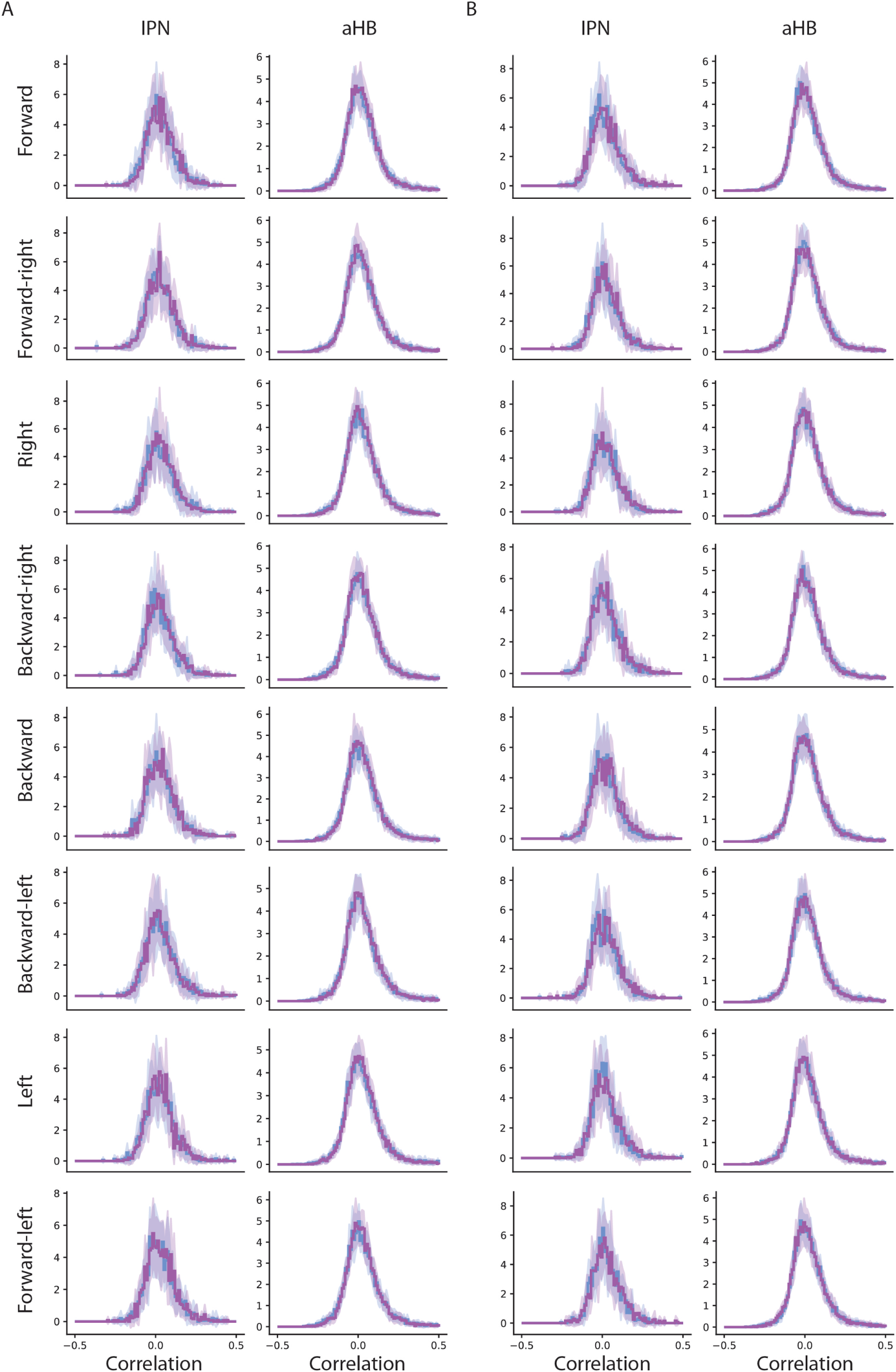
The habenula does not provide visual motion information to the IPN and aHB. (A) distribution of correlation values of IPN and aHB neurons with the visual motion regressors before (blue) and after (purple) NFP treatment for Ntr+ fish (habenula ablated group, mean±sd). The variance of the distribution of correlation values before ablation is not significantly larger than the variance after ablation (p values for each regressor appear in each panel). (B) distribution of correlation values of IPN and aHB neurons with the visual motion regressors before (blue) and after (purple) NFP treatment for control fish (mean±sd). The variance of the distribution of correlation values before ablation is not significantly larger than the variance after ablation (Wilcoxon signed-rank test, p values for all regressor are > 0.99).

**Figure S13:**
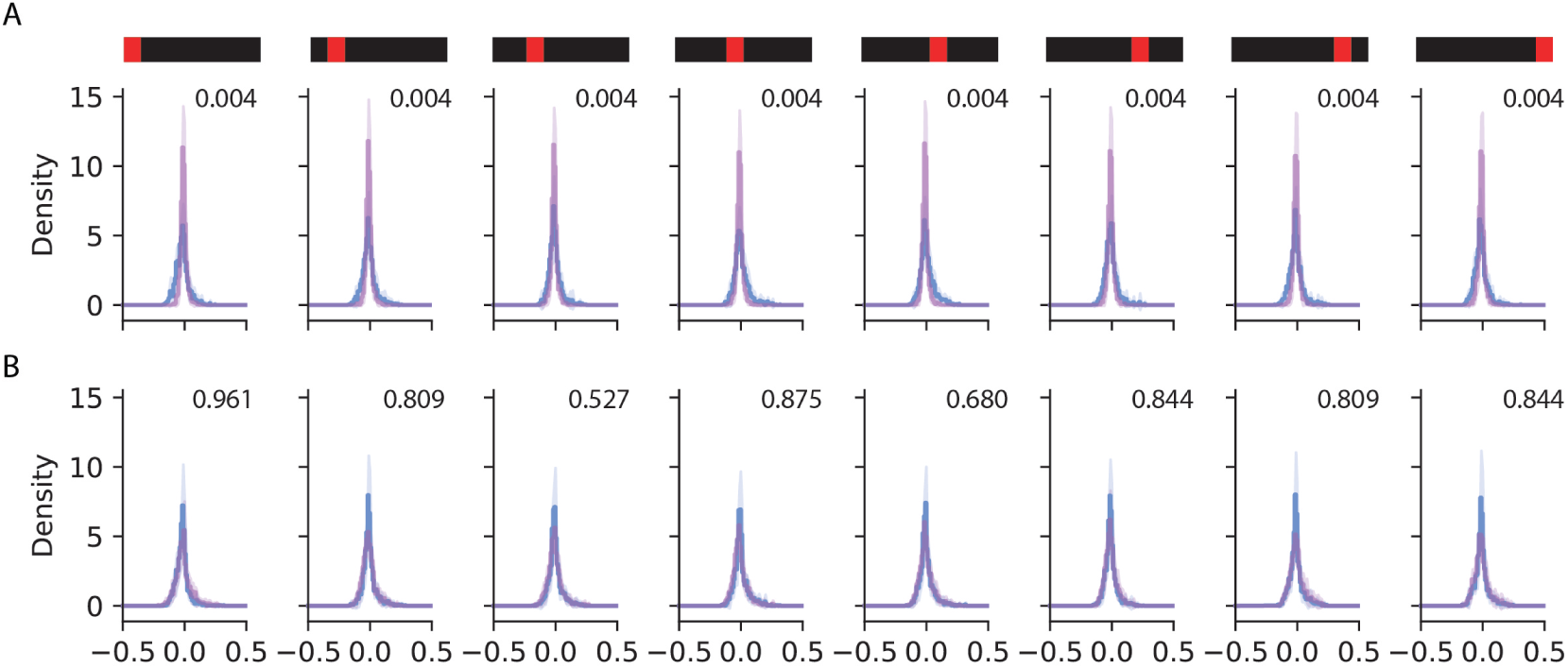
The habenula provides landmark information to the IPN. (A) distribution of correlation values of IPN neurons with the landmark position regressors before (blue) and after (purple) NFP treatment for Ntr+ fish (habenula ablated group, mean ± sd). The variance of the distribution of correlation values before ablation is significantly larger than the variance after ablation (p values for each regressor appear in each panel). (B) distribution of correlation values of IPN neurons with the RF regressors before (blue) and after (purple) NFP treatment for control fish (mean ± sd). The variance of the distribution of correlation values before ablation is not significantly larger than the variance after ablation (Wilcoxon signed-rank test, p values for each regressor appear in each panel).

